# High throughput screen of NPQ in sorghum shows highly polygenic architecture of photoprotection

**DOI:** 10.1101/2025.04.18.649521

**Authors:** Richard L Vath, Samuel B Fernandes, Brandon Monier, Katarzyna Głowacka, Julia Walter, Alexander E. Lipka, John Ferguson, Carl J. Bernacchi, Taylor Pederson, Johannes Kromdijk

## Abstract

- Natural genetic variation in photosynthesis and photoprotection within crop germplasm represents an untapped resource for crop improvement. *Sorghum bicolor* (sorghum) is one of the world’s most widely grown crops, yet the genetic basis of photoprotection in sorghum is not well understood.
- This study examined genetic variation in non-photochemical quenching traits by screening a field-grown panel of 861 genetically diverse natural sorghum accessions across two years.
- Broad-sense heritability ranged between 0.3 to 0.65 across different chlorophyll fluorescence parameters. A combination of genome- and transcriptome-wide (GWAS and TWAS) identification of genetic correlates with the observed trait variation uncovered a complex genetic architecture of many significant small-effect loci. An ensemble approach based on GWAS and TWAS results and the covariance between different fluorescence parameters was used to identify 110 unique candidate genes.
- The resulting high-confidence candidates reveal novel genetic associations with photoprotection and offer resources for further genetic studies and crop genomic improvement efforts.

## Introduction

Growth in global agricultural productivity is likely to be substantially diminished by projected increases in climatic variability (Lewis & King, 2017; Pendergrass et al., 2017; Rahman et al., 2022; Rosenzweig et al., 2014), while demand for food, feed, and biofuel in 2050 is projected to require a 47% increase in agricultural productivity over 2011 levels (Sands et al., 2023). Improvement of crop photosynthetic efficiency has been suggested to offer a potential mitigating solution to help sustain productivity in a changing climate (Zhu et al., 2010). Non-photochemical quenching (NPQ), a leaf physiological process which dissipates excess absorbed light energy (photoprotection), is an important determinant of photosynthetic efficiency, and enhancements in the induction and relaxation rates of NPQ have been shown to increase yield and biomass in C_3_ broad-leaved crops under field conditions (de Souza et al., 2022; Kromdijk et al., 2016). Natural genetic variation in photosynthetic and photoprotective traits promises great potential as a resource for crop improvement (Theeuwen et al., 2022) and variation in these traits has been shown to exist within crop and non-crop germplasm (Ferguson, Caproni, et al., 2023; Kumari et al., 2025; Ortiz et al., 2017; Sahay et al., 2023, 2024; van Rooijen et al., 2017).

*Sorghum bicolor* ((L.) Moench; sorghum) is grown extensively throughout the world as a food and feed crop (Visarada & Aruna, 2019) and offers high potential as a bioenergy stock (Erickson et al., 2012; Rodrigues Castro et al., 2015). Its high water use efficiency and resiliency to stressors (Hadebe et al., 2017; Maman et al., 2003) makes sorghum an attractive option to potentially supplement or replace other crops in increasingly uncertain climate conditions. Sorghum’s adaptability, relatively small diploid genome (∼730 Mb), and extensive genetic diversity facilitate highly tractable variant studies to be conducted in diverse climates. These traits, combined with the availability of a high-quality reference genome (McCormick et al., 2018), make sorghum an attractive resource for understanding genotype-by-environment (GxE) interactions in key C4 photosynthetic traits (Boyles et al., 2019), including NPQ. Uncovering the extent of genetic variability in photoprotective capacity and identifying genes underlying NPQ may help advance breeding efforts toward improvement of photosynthetic efficiency in sorghum.

An understanding of the extent of naturally occurring variation in photoprotection in sorghum, and identification of the genes underlying this variation could facilitate germplasm improvement via breeding, genomic editing, and transgenic approaches (Flood et al., 2011; Lawson et al., 2012; van Bezouw et al., 2019). Genome- and transcriptome-wide association studies (GWAS and TWAS, respectively) can be used to identify quantitative trait loci (QTL) and candidate genes that underlie highly polygenic photosynthetic traits (Gui et al., 2023; Tam et al., 2019; Tibbs Cortes et al., 2021), by correlating genomic marker and transcript expression variation with trait phenotype variation. Despite their demonstrated usefulness, both methods are complicated by the need to set an appropriate threshold for significance, which needs to maintain sufficient stringency to account for the multitude of parallel tests while keeping the rate of false negatives low. However, since the occurrence of false negatives and positives between both methods can be assumed independent, combining GWAS and TWAS offers a way to detect higher-confidence QTL associated with complex crop physiological traits (Ferguson et al., 2021; Kremling et al., 2019; Lin et al., 2022; Pignon et al., 2021).

Molecular and reverse genetics-based approaches have successfully identified genes involved in NPQ regulation in model species (Bru et al., 2020; Kasajima et al., 2011; X.-P. Li et al., 2000), but larger-scale screening of NPQ is a relatively recent pursuit. GWAS have been successful in identifying QTL associated with NPQ in *Arabidopsis thaliana* (Arabidopsis), *Oryza sativa* (rice), *Glycine max* (soybean), *Zea mays* (maize), and sorghum, both in controlled (Rungrat et al., 2019) and field (Ferguson, Caproni, et al., 2023; Herritt et al., 2016; Kumari et al., 2025; Sahay et al., 2023, 2024; Wang et al., 2017) conditions. Screening of NPQ kinetics on field-grown plants can be cumbersome and challenging to accomplish in a manner which is high-throughput enough for the large number of accessions required to discern small-effect loci via GWAS and TWAS. As a result, our understanding of the genes behind natural variation in NPQ under production-relevant conditions is still limited for several of the world’s most important crops, particularly those with C_4_ photosynthetic metabolism, such as maize, *Saccharum officinarum* (sugarcane), and sorghum.

The current study utilises a high-throughput chlorophyll fluorescence screening method (Ferguson et al., 2023a, 2023b; Gotarkar et al., 2022) to characterise and quantify rates of NPQ induction and relaxation in field-grown plants of 861 biomass sorghum accessions across two years. Most of the measured traits showed moderately high broad-sense heritability and their observed variation was underpinned by a complex architecture of many significant small-effect loci. Combined GWAS and TWAS analyses uncovered 110 unique high-confidence candidate genes, which may be used to improve understanding of NPQ regulatory regions in sorghum and other economically important, closely-related C4 crops such as maize and sugarcane, and potentially in even more distantly related species (Sahay et al., 2023, 2024).

## Materials and Methods

### Germplasm and field trial design

A sorghum panel composed of 869 genetically diverse biomass accessions (described previously in dos Santos et al., 2020; Valluru et al., 2019)) was grown in 2017 and 2019 at the University of Illinois Maxwell Farm (2017; 40.055166, -88.236794) and Energy Farm (2019; 40.063333, - 88.205000) Research Sites near Urbana, IL, USA. Accessions were planted in an augmented block design with 960 four-row plots (3 m row length) arranged into 40 columns and 24 rows, in 16 blocks. The incomplete blocks were connected through six common check accessions to account for block effects during statistical analysis of phenotypic traits. This study utilised data from 855 and 846 accessions in 2017 and 2019, respectively, after filtering for availability of marker data. Together this allowed for a combined-year analysis of 861 accessions. 839 accessions were common to both years, with three check accessions (Pacesetter, PI276801, and PI148089) common between years. The panel was planted on May 31 of both years, with five plots replanted on June 10, 2019 due to poor germination. Temperature and precipitation data for both growing seasons are available in Supporting information Figure S1.

### Field sampling

Sorghum accessions were screened for photoprotective traits in both years via chlorophyll fluorescence imaging. Sampling and screening were conducted using detached leaf segments (Ferguson et al., 2023b) via the 96-well plate method detailed in Gotarkar et al. (2022) and Sahay et al. (2023). Plants were sampled by cutting leaf discs with a 6 mm diameter hole punch from the youngest fully expanded leaf, as indicated by ligule emergence at the time of measurement. Two samples from separate plants in the middle of both inner rows from each plot were taken, providing a total of four biological replicates per accession. To prevent sampling time differences between accessions, blocks were sampled sequentially by replicate. Hence, sampling time varied between replicates, but differences in average sampling time between genotypes were kept within 45 min (time taken to sample one block). Following collection, sample plates were wrapped in aluminium foil to prevent light intrusion and buffer temperature changes, then stored in a cooled polystyrene container while further sampling was completed. Four replicates of 128 accessions were sampled per day, between 14:30 and 18:30 local time. After all daily sampling was completed, plates were stored overnight at approximately 20°C in a temperature-controlled lab. Sampling took place in 2017 from July 25 to 28 and August 1 to 4, and in 2019 from July 22 to 25 and July 28 to 31.

### Chlorophyll fluorescence screening of photoprotective traits

Discs were imaged on the morning proceeding sampling using a fluorescence imager (CFImager, Technologica, Colchester, UK). The 96-well plates were imaged in the order in which they were sampled the previous afternoon, to mitigate temporal bias. In a dimly lit room, foil was removed from each plate immediately before imaging, and the plate placed into the imager. The imaging routine utilised 800 ms long flashes at 4,000 μmol m^-2^ s^-1^ photosynthetic photon flux density (PPFD), and consisted of a dark-adapted measurement of maximum photosystem II (PSII) quantum efficiency (*Fv*/*Fm*) followed by periodic fluorescence measurements over 10 minutes at 2,000 μmol m^-2^ s^-1^ actinic PPFD followed by 12 minutes of darkness. During the NPQ induction (light) period, fluorescence parameters were measured every 20 seconds for the first minute, then every minute for the remainder of the light period. During the NPQ relaxation (dark) period, fluorescence parameters were measured every 20 seconds for the first minute, then every minute for the next three minutes, then every three minutes for the duration of the dark period (Figure 1). Image thresholding and segmentation were performed in MatLab (*MATLAB*, 2020). Fluorescence values for each disc were taken as the median pixel value of the disc. The distribution of *Fv*/*Fm* values for the discs was examined to discern outlier leaf samples and discs with *Fv*/*Fm* values lower than 0.65 were excluded from further analyses.

**Figure 1:**
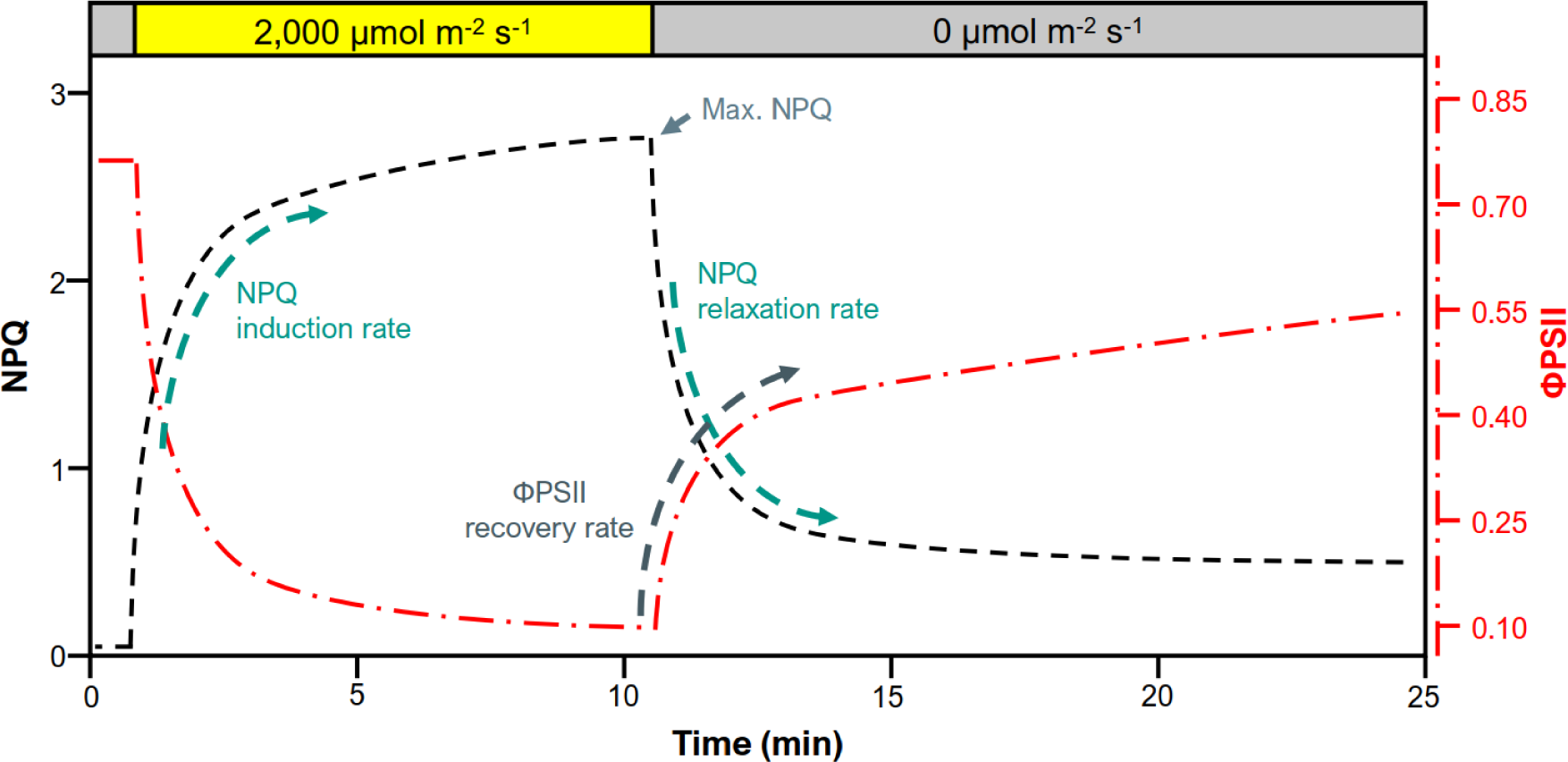
Illustrative plot of NPQ induction during a high-light treatment followed by NPQ relaxation and 𝜱*PSII* recovery during a subsequent period of darkness (styled after Kaiser et al., 2018).

Using MatLab, exponential models were fit to the NPQ traces of light induction and dark relaxation periods separately, to allow quantitative comparison of NPQ induction (Eq. 1) and relaxation (Eq. 2) rates, as well as the dark recovery of PSII operating efficiency (𝛷*PSII*, Eq. 3):

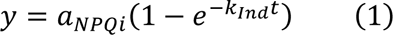

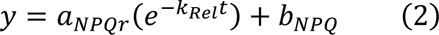

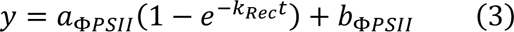

where 𝑎_*NPQi*_, 𝑎_*NPQr*_and 𝑎_*𝛷PSII*_are the initial values of NPQ induction, relaxation, and *𝛷PSII* recovery during the dark period, respectively; *K_Ind_*, *K_Rel_*, and *K_Rec_* the rate constants of NPQ induction and relaxation and *𝛷PSII* recovery, respectively (thus a larger number indicates faster kinetics), *t* is the measurement time point, and *b_NPQ_* or *b_ΦPSII_* an offset to account for a non-zero y-intercept term. Modelled curve fits were visually examined for goodness-of-fit, with all traits of non-conforming discs excluded from further analyses, resulting in the exclusion of 8% of discs in 2017 and 13% in 2019. Figure 1 provides a stylised depiction of fluorescence trace parameters.

In addition to the parameters derived from the exponential models, maximum NPQ reached during the light period and the initial linear slopes of NPQ induction/relaxation were also determined for each trace and the photoprotection index (𝑃𝐼), depicting the photoprotective effectiveness of NPQ, was calculated based on the method described by Ruban & Murchie (2012) and implemented as in Kromdijk et al. (2016). Briefly, 𝑃𝐼 is the ratio between observed *Fv*′/*Fm*′ (maximum PSII quantum efficiency at a given PPFD) and the calculated final *Fv*′/*Fm*′ based on the predictable decrease due to residual NPQ by the end of the 12-minute dark period. As in Ruban & Murchie (2012), 𝑃𝐼 ≥ 1 suggests that the entire reduction in *Fv*′/*Fm*′ relative to initial *Fv*/*Fm* can be attributed to NPQ, while values progressively lower than one suggest a portion of the drop is attributable to photoinhibition (sustained depression of *Fv*′/*Fm*′_𝑓_ due to reaction centre damage). 𝑃𝐼 was calculated as follows:

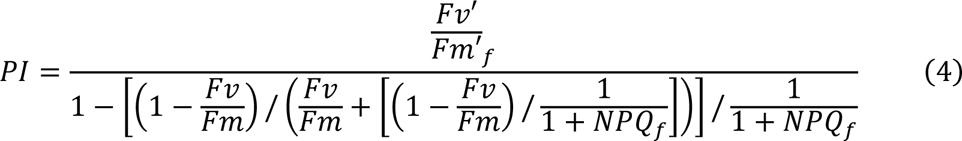

where *Fv*′/*Fm*′_𝑓_ and 𝑁𝑃𝑄_𝑓_ are *Fv*′/*Fm*′ and 𝑁𝑃𝑄, respectively, at the final dark time point.

### Statistical modelling and heritability

For each trait/year combination a residual maximum likelihood model was fit to each trait using ASReml for R (Butler et al., 2017):

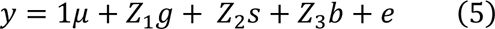

where 𝑦 is the vector of phenotypes; 1 (𝑛 x 1) is a vector of ones; μ is the trait mean; 𝑍_1_ is the incidence matrix associated with the vector of random genotype (accession) effects 𝑔, with 𝑔 ∼ 𝑁(0, 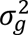) where 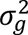 is the genetic variance; *Z*_2_ is the incidence matrix associated with the vector of random effect set 𝑠, with 𝑠 ∼ 𝑁(0, 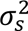) where 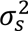 is the variance of set; *Z*_3_ is the incidence matrix associated with the vector of random effect block within set 𝑏, with 𝑏 ∼ 𝑁(0, 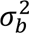) where 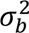 is the variance of block within set; and 𝑒 is the vector of residuals, with 𝑒 ∼ 𝑁(0, 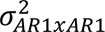) where 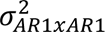 is the residual variance with a first-order auto-regressive structure applied to row and column for spatial correction.

Additionally, the two years’ data were analysed in a joint model:

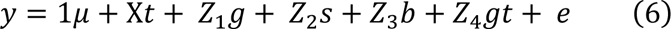

where 𝑦 is the vector of phenotypes for j environments; 1 (𝑛 x 1) is a vector of ones; μ is the trait mean; 𝑋 is the incidence matrix associated with the vector of fixed effect environments 𝑡 (j x 1); 𝑍_1_ is the incidence matrix associated with the vector of random genotype effects within environment 𝑔, with 𝑔 ∼ 𝑁(0, 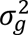) where 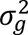 is the genetic variance; 𝑍_2_ is the incidence matrix associated with the vector of random effect of set within environment 𝑠, with 𝑠 ∼ 𝑁(0, 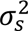) where 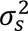 is the variance of set within environment; 𝑍_4_ is the incidence matrix associated with the vector of random effect block within set within environment 𝑏 , with 𝑏 ∼ 𝑁(0, 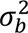), where 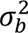 is the variance of block within set within environment; 𝑍_4_ is the incidence matrix associated with the vector of random effect genotype interacting with environment 𝑔𝑡, with 𝑏 ∼ 𝑁(0, 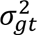), where 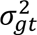 is the variance of genotype with environment; and 𝑒 is the vector of residuals, with 𝑒 ∼ 𝑁(0, 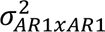) where 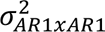 is the residual variance with a first-order auto-regressive structure applied to row and column for spatial correction. The most appropriate variance-covariance structure to model the residuals was selected based on the Akaike information criterion. Outliers were filtered out based on method two of Bernal-Vasquez et al. (2016).

Best linear unbiased predictions (BLUPs) were obtained separately for each genotype for 2017, 2019, and the combined model, resulting in data for 861 accessions to be used for genomic analysis. Generalised heritability (analogous to broad-sense heritability) for each trait was calculated as

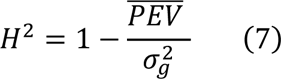

where PEV is the prediction error variance (Cullis et al., 2006; Piepho & Möhring, 2007) and 𝜎_𝑔_ is genetic variance.

Correlations between BLUPs were calculated and visualised with R packages *psych* (Revelle, 2022) and *ggcorrplot2* (Cai et al., 2022).

### Genome-wide association

The genotype data set utilised in this study was previously published in Ferguson et al. (2021). Briefly, 100,435 genotyping by sequencing single nucleotide polymorphisms (SNPs) available for 869 accessions (dos Santos et al., 2020) were imputed using a whole genome re-sequencing panel with 5,512,653 SNPs and 229 accessions from Valluru et al. (2019). Imputation was done with Beagle 4.1 (Browning & Browning, 2016) after filtering out SNPs with a minor allele count less than 20 and pruning SNPs in high linkage disequilibrium (LD) (r^2^ > 0.9), using Plink (Purcell et al., 2007) with options “–indep-pairwise 50 10 0.9”. The resulting data sets had 450,074 (2017), 450,449 (2019), and 454,087 (joint analysis) SNPs that were used for GWAS. The SNP dataset was used in TASSEL 5 (Bradbury et al., 2007) to obtain the kinship matrix and five principal components (PCs). In both cases, the default options were used. Univariate and multivariate GWAS were conducted in GEMMA (Zhou & Stephens, 2012) using the Q+K model for each trait/year combination. The best number of principal components was decided based on the Bayesian information criterion.

Analysis of specific combinations of covarying traits might increase signal to noise for specific NPQ attributes of interest and potentially suggest genotypes which are particularly and uniquely responsive to photoprotection under dynamic light conditions. Thus, GWAS was performed on several sets of combined traits (CT). For multi-trait GWAS, the following trait combinations were evaluated:

- CT1: Max NPQ, NPQ induction amplitude, NPQ induction *k*, NPQ relaxation *k*
- CT2: NPQ induction amplitude, NPQ induction *k*, NPQ relaxation *k*
- CT3: Max NPQ, NPQ induction *k*, NPQ relaxation *k*
- CT4: 𝑃𝐼, 𝛷𝑃𝑆𝐼𝐼 recovery amplitude, 𝛷𝑃𝑆𝐼𝐼 recovery 𝑘

CT1 through CT3 are expected to amplify signal along the quickly vs. slowly relaxing NPQ axis of variation, while CT4 should amplify variation along the photoprotection vs. photodamage axis.

The unpruned dataset was used to calculate pairwise linkage LD, only including SNPs with an r^2^ above 0.2, and using a sliding window maximum size of 500kb and 99,999 SNPs with Plink options “–blocks no-pheno-req no-small-max-span, –blocks-max-kb 500, –blocks-min-maf 0.001”. LD blocks were calculated as proposed in Gabriel et al. (2002). SNPs were considered to be in strong LD if the bottom 90% D-prime confidence interval was greater than 0.70, and the top of the confidence interval was at least 0.98, for a total of 45,311 LD blocks (see Supporting information Table S1) with a median size of 1.178kb containing a median of 11 SNPs. Manhattan plots for visualisation of SNP mapping were based on the “myManhattan” function (https://github.com/alfonsosaera/myManhattan) with modifications.

### Transcriptome-wide association

Gene expression data (described in Ferguson et al. (2021)) from the controlled environment-grown 229 sorghum accessions of Valluru et al. (2019) was used to analyse covariance of transcript abundance with photoprotection traits. Transcript abundance from tissue at the shoot growing point (GP) and base of the third-leaf (3L) was analysed separately for each photoprotection trait/year combination.

Before mapping, ten hidden factors were calculated using probabilistic estimation of expression residual (PEER) factors for each tissue (Stegle et al., 2012). Five PCs were also calculated from prior genotype data. Finally, genes expressed in less than half of the individual lines were removed from each tissue set. After covariate calculation and filtering, a general linear model was fit individually for each NPQ trait and gene expression value using prior PEER factors and PCs as covariates. In addition to mapping each NPQ trait, a multitrait approach was also performed by combining several traits using methods described in the prior section. Mapping was conducted in the R environment (R Core Team, 2022) using *rTASSEL* (Monier et al., 2022).

### Combined genome and transcriptome-wide analysis

Fisher’s combined test (FCT) was performed using the GWAS and TWAS results to reduce the impact of false positives and negatives, based on the fact that the likelihood of true functional genetic variation increases dramatically when multiple types of analyses are performed and co-analysed (Kremling et al., 2019). For each trait/year combination, the nearest gene (by physical location) was assigned to the top 10% of GWAS SNPs by p-value. FCT was then run on the combined top GWAS and all TWAS transcripts using the *metap* package for R (Dewey, 2022).

### Candidate gene selection and investigation

Candidate genes (Supporting information Table S2) were selected utilising an ensemble approach (Kremling et al., 2019) which increases the statistical power available for determining genes associated with leaf physiological traits, which are often highly polygenic. Genes within LD blocks of the top 0.05% of GWAS SNPs (Supporting information Table S3) and the top 1% of genes from each TWAS (Supporting information Table S4) and FCT analysis (Supporting information Table S5), by *p*-value, were classified as “top genes”. Fifteen total analyses were performed for each trait, comprising of 2017, 2019, and joint models for GWAS and 2017, 2019, and joint models for both TWAS and FCT based on GP and 3L tissue.

Arabidopsis orthologues of sorghum genes overlapping in more than three separate analyses were identified using R package *gProfiler2* (Kolberg et al., 2020). Panther (http://pantherdb.org/) statistical over-representation analysis for biological function was performed on the Arabidopsis gene list against the Arabidopsis reference genome via *rbioapi* (Rezwani et al., 2022) (Supporting information Table S6).

Top sorghum genes were considered candidates for photoprotection based on: 1), identification in eight or more individual analyses for a given trait, and 2) identification as a top-ranked gene in 10 or more separate traits (Figure 2; Supporting information Table S7). Eight analyses and ten traits were chosen as thresholds because the number of candidate genes overlapping multiple analyses or traits grew exponentially larger when fewer analyses or trait overlaps were considered. Additionally, several top sorghum genes were considered candidates based on manual investigation via direct searches for the sorghum gene ID in Google Scholar and based on their corresponding Arabidopsis orthologue being annotated in TAIR (Berardini et al., 2015) for light response and photoprotection related traits (such as xanthophyll synthesis or nonphotochemical quenching).

**Figure 2:**
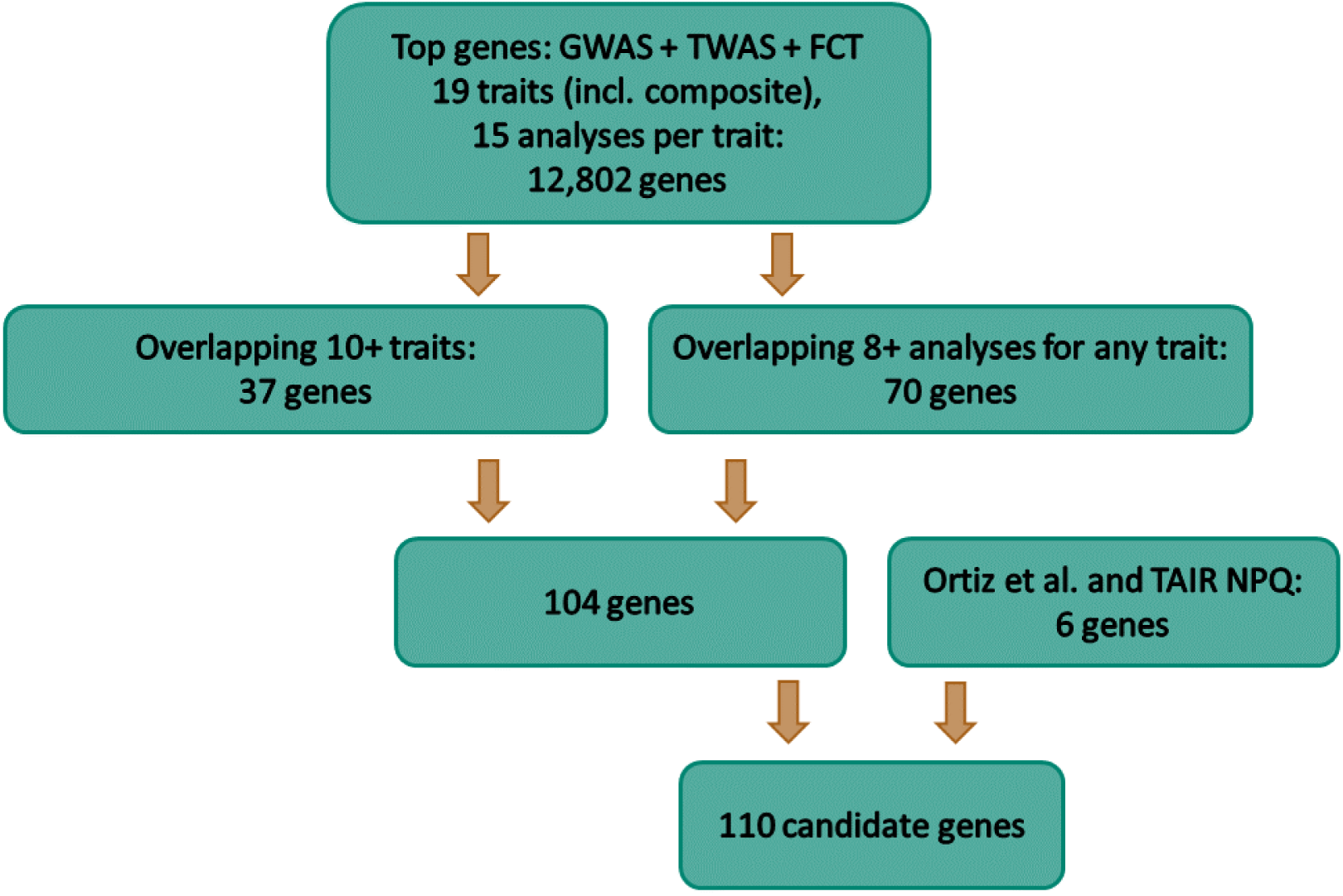
Workflow of sorghum candidate gene selection based on top 0.05% of GWAS and top 1% of TWAS and FCT genes overlapping multiple traits and analyses, and based on manual investigation.

Sorting intolerant from tolerant (SIFT) analysis (Ng & Henikoff, 2003) was performed on candidate genes to investigate the likelihood of coding-region SNPs’ contribution to variation in protein function, utilising SIFT scores from a sorghum panel containing 286 of the accessions used in this study (Lozano et al., 2021). Upset plots (Lex et al., 2014) were produced using the *UpSetR* package (Conway et al., 2017) to aid in visualisation of top gene set overlaps for individual traits.

## Results

### Genetic diversity in sorghum harbours substantial variation in photoprotection traits

A high-throughput chlorophyll fluorescence screening method was employed to characterise photoprotective trait diversity in a field-grown sorghum panel. NPQ induction and relaxation kinetics were measured in each sorghum accession via the application of a high-light treatment followed by a dark period, allowing for a quantitative comparison of photoprotective capacity across a genetically diverse sorghum population. Substantial variation was observed among sorghum accessions in NPQ traits. The percentage difference between the lowest and highest accessions in maximum NPQ, NPQ induction rate constant, NPQ relaxation rate constant, and rate constant of 𝛷𝑃𝑆𝐼𝐼 recovery were 24%, 115%, 87%, and 63%, respectively, for joint model BLUPs of each trait (Table 1 and Figure 3).

**Figure 3:**
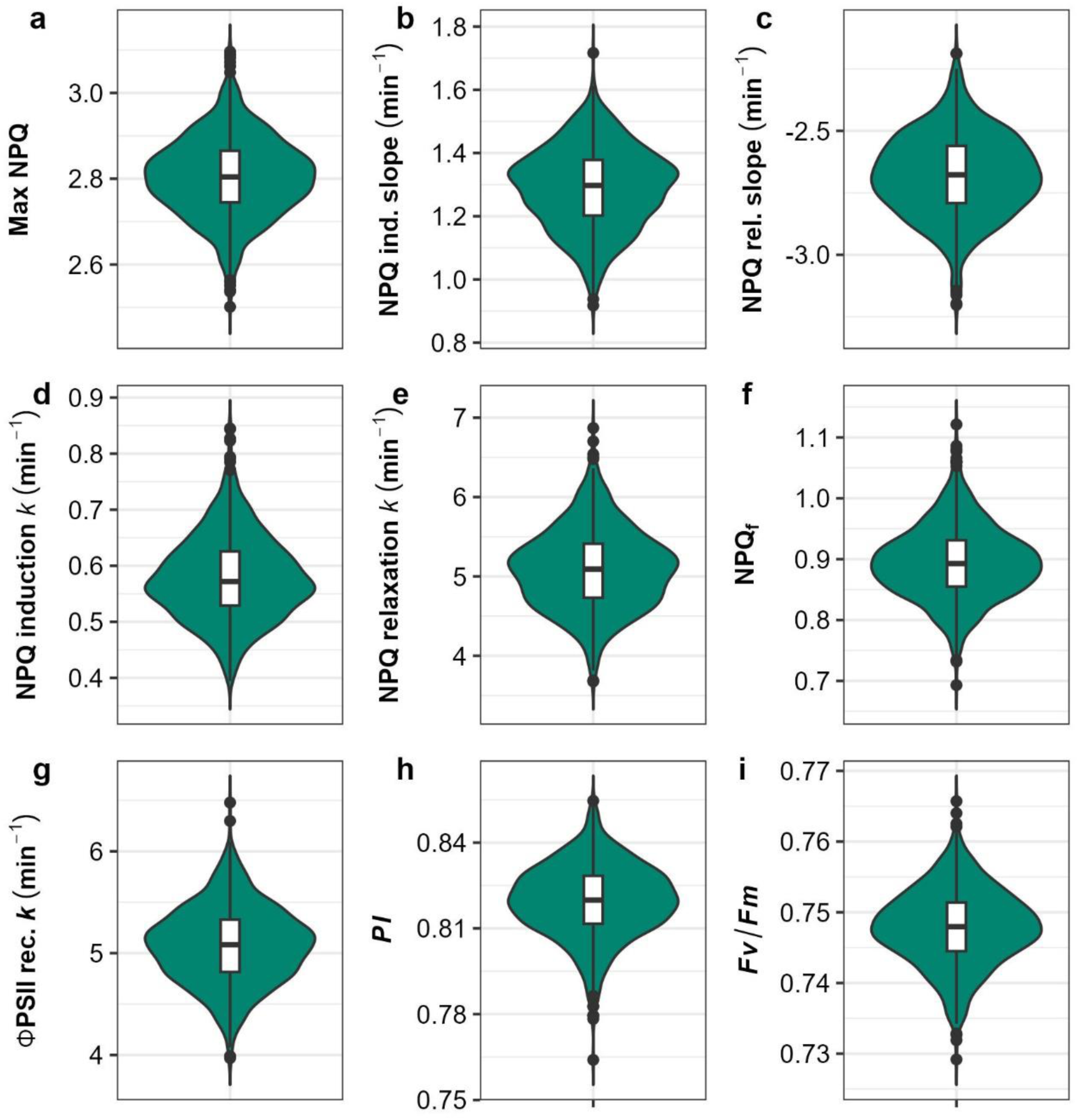
Violin plots of variation in adjusted genotype means of NPQ trace parameters for the joint model. Internal box plot edges represent first and third quartiles. Points represent outliers beyond 1.5 times the interquartile range. The solid line within the boxes indicates the median. a, maximum NPQ level reached during the trace; b, linear slope of NPQ induction; c, linear slope of NPQ relaxation; d, rate constant *k* of NPQ induction; e, rate constant *k* of NPQ relaxation; f, residual NPQ at end of dark relaxation period; g, rate constant *k* of 𝜱*PSII* recovery; h, photoprotection index; i, 𝑭𝒗/𝑭𝒎.

**Table 1:** Descriptive statistics of photoprotection traits for joint adjusted means (BLUPs) of 861 sorghum accessions.

| Trait | Description | Mean | Min | Max | $H^2$ |
| --- | --- | --- | --- | --- | --- |
| Max NPQ | Maximum NPQ level reached during fluorescence trace | 2.80 | 2.50 | 3.10 | 0.52 |
| NPQ ind. slope | Initial linear slope of NPQ induction ($\text{min}^{-1}$) | 1.29 | 0.92 | 1.72 | 0.58 |
| NPQ rel. slope | Initial linear slope of NPQ relaxation in dark ($\text{min}^{-1}$) | -2.68 | -3.20 | -2.19 | 0.67 |
| NPQ ind. $k$ | Exponential rate constant of NPQ induction ($\text{min}^{-1}$) | 0.58 | 0.39 | 0.84 | 0.67 |
| NPQ rel. $k$ | Exponential rate constant of NPQ relaxation in dark ($\text{min}^{-1}$) | 5.09 | 3.68 | 6.87 | 0.44 |
| $NPQ_f$ | NPQ at final dark time point | 0.89 | 0.69 | 1.12 | 0.47 |
| $\Phi PSII$ rec. $k$ | Exponential rate constant of $\Phi PSII$ recovery in dark ($\text{min}^{-1}$) | 5.08 | 3.97 | 6.48 | 0.43 |
| $PI$ | Photoprotection index | 0.82 | 0.76 | 0.85 | 0.36 |
| $Fv/Fm$ | Maximum $PSII$ quantum efficiency | 0.75 | 0.73 | 0.77 | 0.30 |

331 *Note:* Trait calculations detailed in *methods* and illustrated in Figure 1. $H^2$ is heritability

While the range of maximum NPQ remained similar in both 2017 and 2019, NPQ kinetic and light-use efficiency traits varied considerably over the two years (Figure 4), potentially due to weather differences between both years (Supporting information Figure S1) with total precipitation was substantially higher in 2019. NPQ induction in 2019 was slower across the panel, with a slope of 1.19 min^-1^ and *k* of 0.53 min^-1^, compared to a slope of 1.41 min^-1^ and *k* of 0.62 min^-1^ in 2017. NPQ relaxation was faster in 2019, with a median slope of -2.91 min^-1^ and *k* of 6.19 min^-1^, compared to a slope of -2.44 min^-1^ and *k* of 3.97 min^-1^ in 2017. The rate of 𝛷𝑃𝑆𝐼𝐼 recovery was also faster in 2019, with a median value of 5.66 min^-1^ compared to 4.47 min^-1^ in 2017. These differences may relate to inter-year variation in susceptibility to photoinhibition during the light treatment, as the median 𝑃𝐼 in 2019 was 0.85 compared to 0.79 in 2017. Pairwise scatterplots of 2017 and 2019 BLUPs are shown in Figure 5. Accession values for all traits were weakly to moderately correlated between years with Pearson’s correlation (*r*) values ranging between 0.21 and 0.45, but correlations were strongly significant in all cases, suggesting that accession ranks were generally maintained despite fairly strong genotype-environment interaction effects.

**Figure 4:**
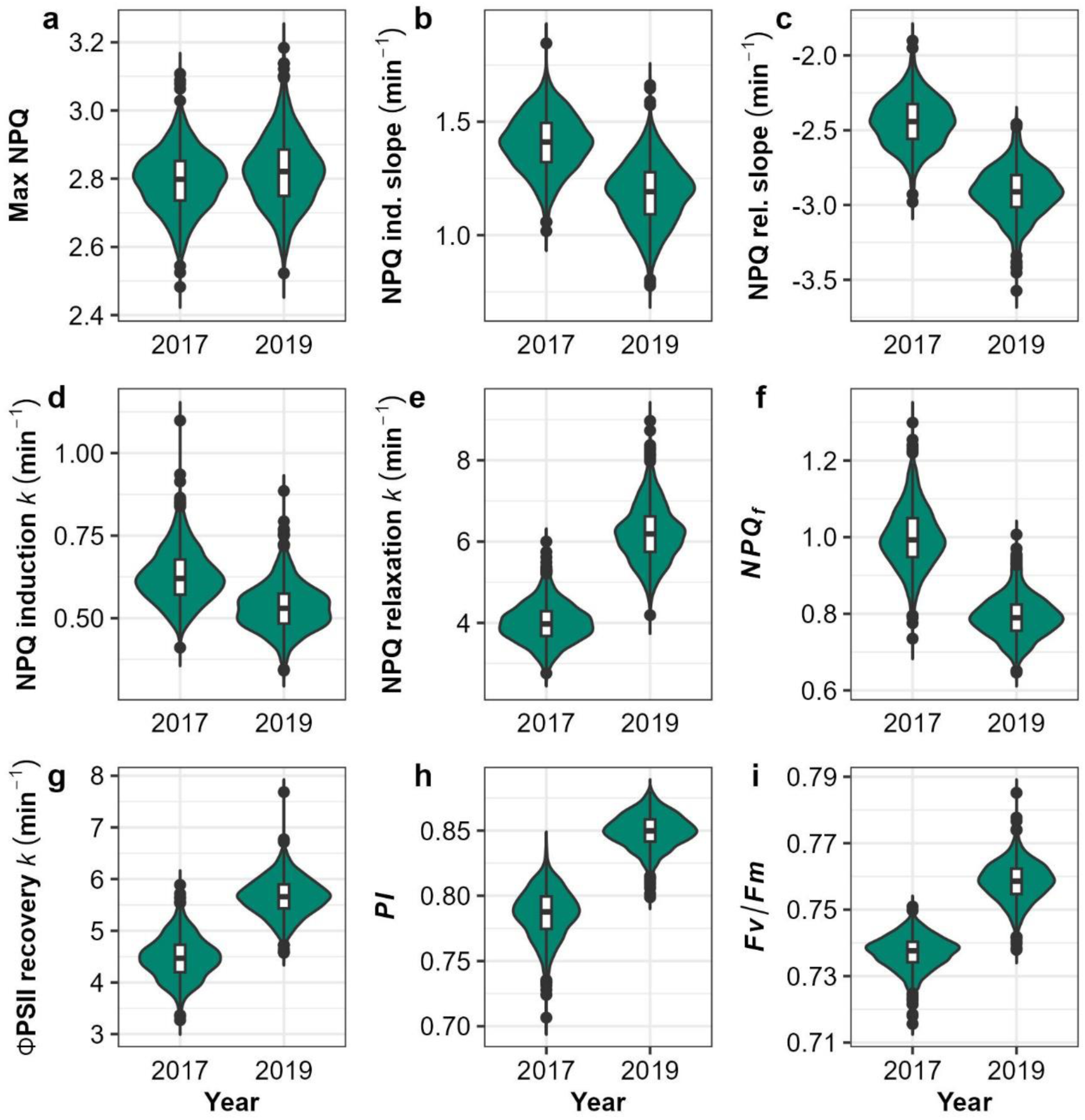
Violin plots of variation in adjusted genotype means of NPQ trace parameters for 2017 and 2019 field seasons. Internal box plot edges represent first and third quartiles. Points represent outliers beyond 1.5 times the interquartile range. The solid line within the boxes indicates the median. a, maximum NPQ level reached during the trace; a, linear slope of NPQ induction; c, linear slope of NPQ relaxation; d, rate constant *k* of NPQ induction; e, rate constant *k* of NPQ relaxation; f, residual NPQ at end of dark relaxation period; g, rate constant *k* of 𝜱*PSII* recovery; h, photoprotection index; i, 𝑭𝒗/𝑭𝒎.

**Figure 5:**
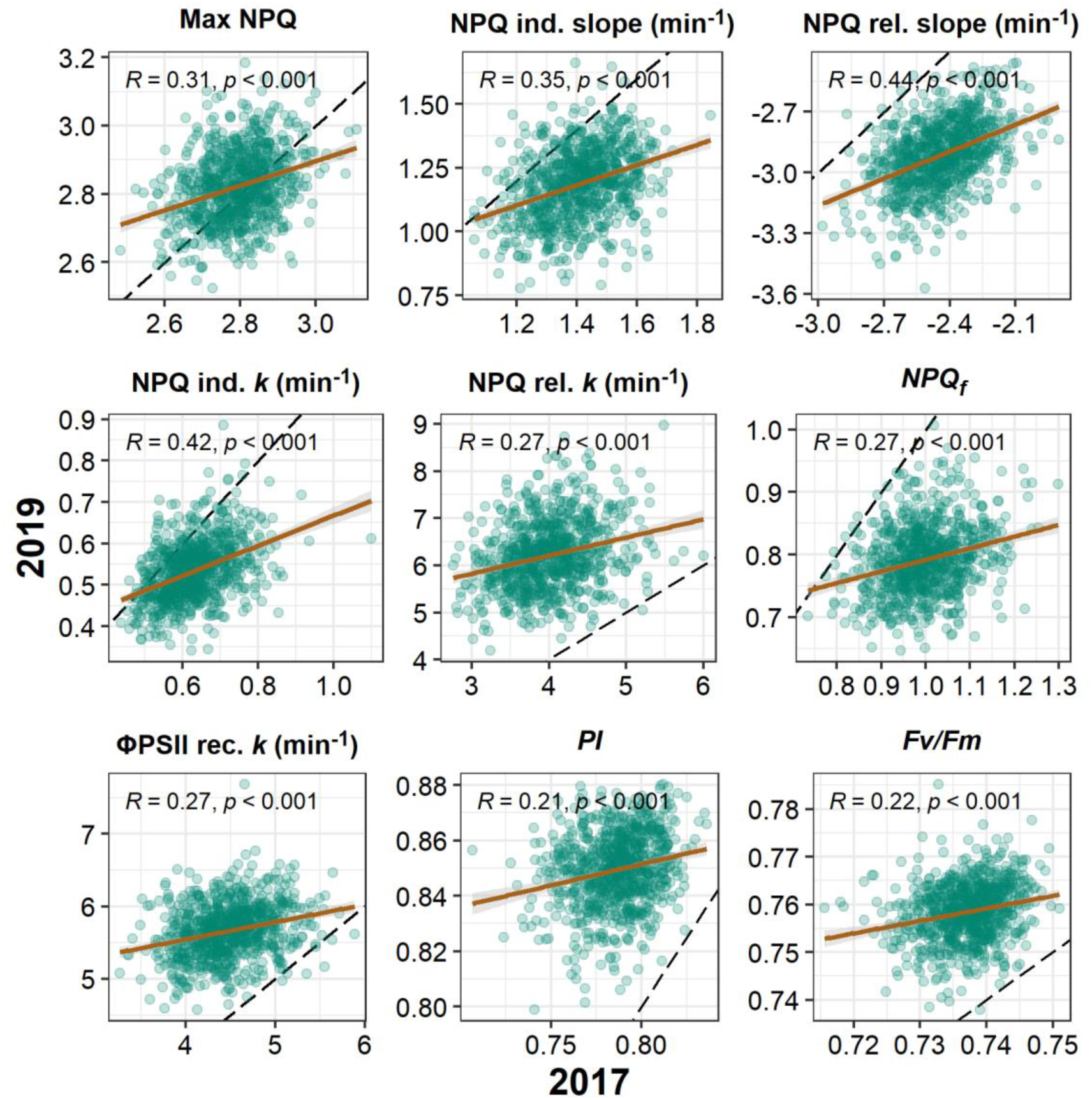
Relationships of 2017 and 2019 BLUPs including Pearson’s *r* and *p*-value. Linear regression line shown in brown. The dashed black line is the 1:1 line.

Heritability (H^2^) of NPQ traits was moderate to moderately high (Figure 6, with joint model H^2^ for maximum NPQ at 0.52, NPQ induction and relaxation slopes at 0.58 and 0.67, respectively, and NPQ induction and relaxation *k* at 0.67 and 0.44, respectively. Light use efficiency traits and 𝑃𝐼 were less heritable, with joint BLUP H^2^ values of 0.43 and 0.30 for 𝛷𝑃𝑆𝐼𝐼 recovery *k* and *Fv*/*Fm*, respectively, and 0.36 for 𝑃𝐼. Values of H^2^ for 2017 and 2019 models were consistent across years for most traits, excepting 𝑃𝐼 and *Fv*/*Fm* which were notably higher in 2019 (0.60 for 𝑃𝐼 and 0.45 for *Fv*/*Fm*) than in 2017 (0.47 for 𝑃𝐼 and 0.37 for *Fv*/*Fm*).

**Figure 6:**
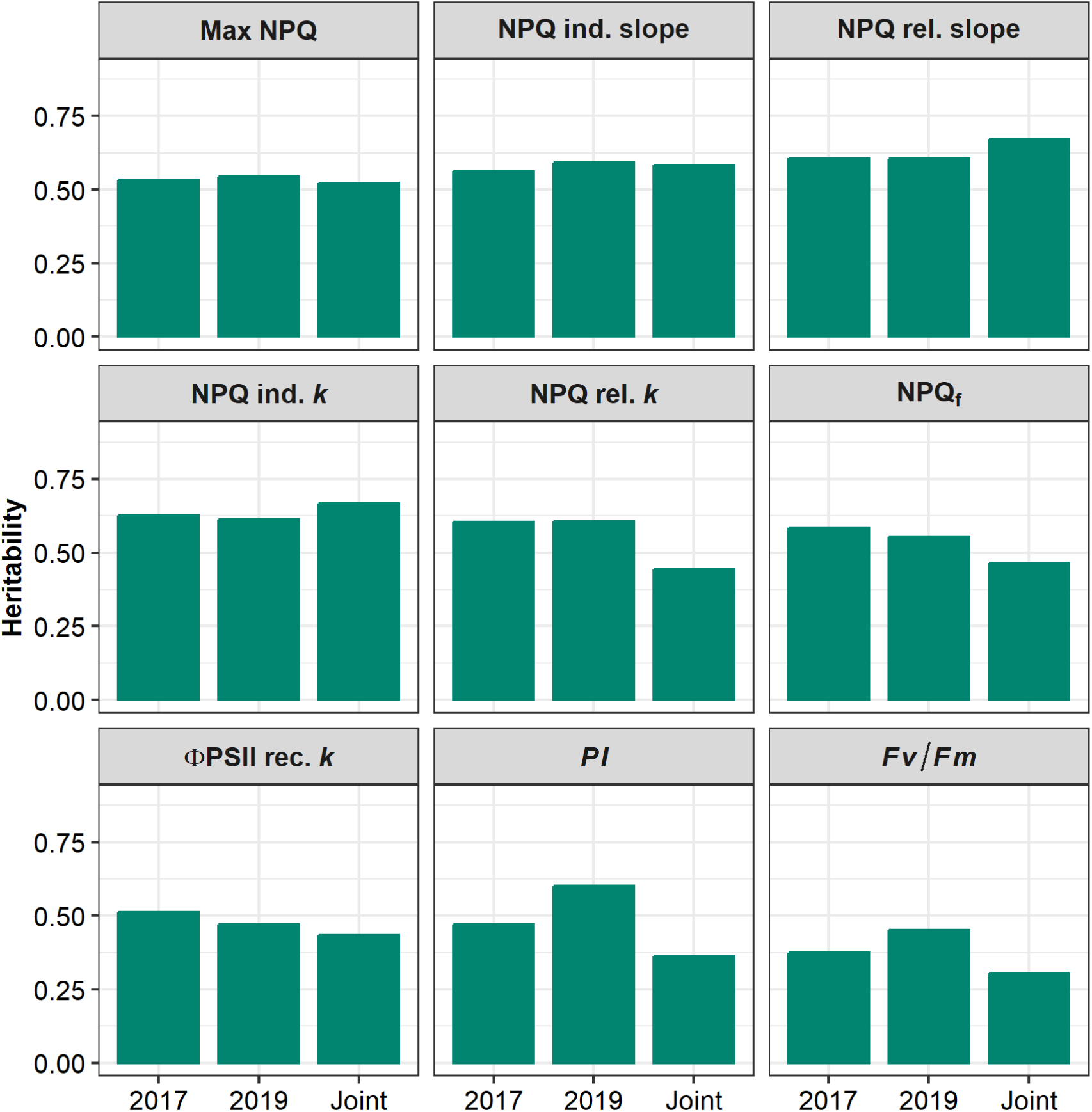
Bar plots of heritabilities of sorghum photoprotective traits calculated from 2017, 2019, and joint analysis BLUPs. Max NPQ, maximum NPQ level reached during the trace; NPQ ind. / rel. slopes, initial linear slopes of NPQ induction and relaxation, respectively; NPQ ind./ rel. *k*, exponential rate constants of NPQ light induction and dark relaxation, respectively; NPQ_f_, NPQ at final dark time point; 𝜱*PSII* rec. *k*, Exponential rate constant of 𝜱*PSII* recovery in dark; 𝑷𝑰 , photoprotective index; 𝑭𝒗/𝑭𝒎 , Maximum PSII quantum efficiency.

Correlations between traits were consistent between BLUPs estimated for each year (Figure 7 **b & c**) and between joint model BLUPs (Figure 7 **a**). Maximum NPQ tended to correlate with slower NPQ induction and faster relaxation kinetics: Significant (*p* < 0.05) correlations were observed between max NPQ and NPQ induction *k* (Pearson’s *r* = -0.33, joint model), initial relaxation slope (*r* = -0.74; joint model), and relaxation *k* (*r* = 0.13; joint model). NPQ relaxation *k* was also strongly positively correlated with 𝜱PSII recovery *k* (*r* = 0.85; joint model), suggesting the short high light treatment did not cause substantial reaction centre damage which might otherwise disrupt the expected strong relationship between 𝜱PSII and NPQ. As could be expected, initial slopes of NPQ induction and relaxation were significantly correlated with their corresponding rate constants *k*, at *r* = 0.88 for induction and *r* = -0.48 for relaxation (joint model). Intriguingly, accessions with a higher NPQ induction *k* tended to have a slower (less negative) initial relaxation slope (*r* = 0.25), both of which would enhance protection against photoinhibition. Accessions with lower 𝑷𝑰 (more photoinhibited) tended to exhibit a higher 𝑵𝑷𝑸_𝒇_ (*r* = -0.72), suggesting that the final NPQ level determined at the end of the imaging assay also reflects a degree of photoinhibition sustained during the light treatment.

**Figure 7:**
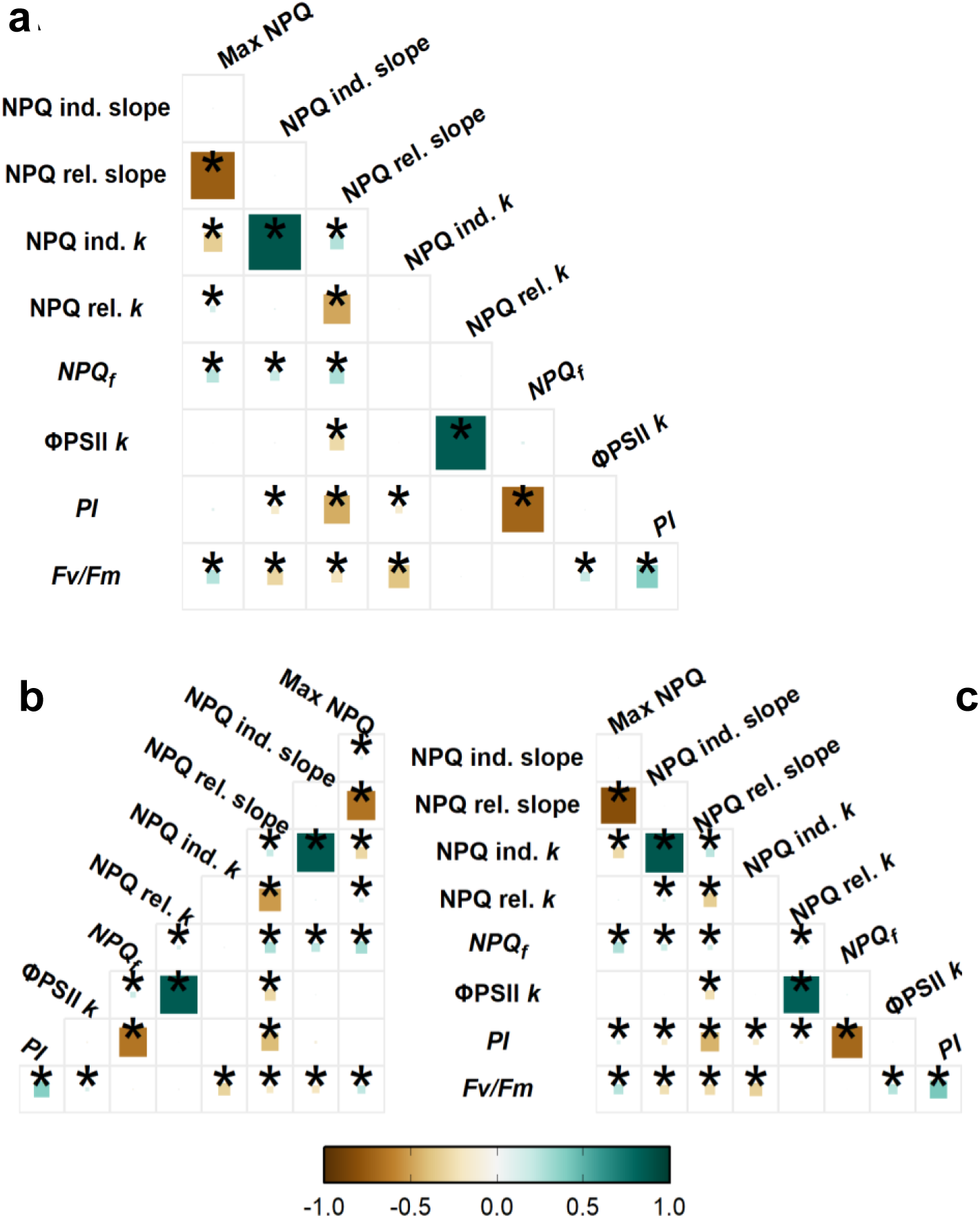
Correlograms demonstrating Pearson’s correlations between BLUPs of photoprotection traits of 839 sorghum accessions. The colour and size of e ach square represent the correlation coefficient of the pairwise interaction. Squares marked with an asterisk denote Holm-method corrected *p*-values < 0.05. A, joint model. B, 2017 model. C, 2019 model. Max NPQ, maximum NPQ level reached during the trace; NPQ ind./rel. slopes, initial linear slopes of NPQ induction and relaxation, respectively; NPQ ind./ rel. *k*, exponential rate constants of NPQ light induction and dark relaxation, respectively; NPQf, NPQ at final dark time point; 𝜱*PSII* rec. *k*, Exponential rate constant of 𝜱*PSII* recovery in dark; 𝑷𝑰, photoprotective index; 𝑭𝒗/𝑭𝒎, Maximum *PSII* quantum efficiency.

### Genome- and transcriptome-wide analyses uncover genes associated with photoprotection in sorghum

Marker-trait association (MTA) analyses were performed using the BLUPs from 2017, 2019, and combined year (joint) models, to identify sorghum genomic regions associated with variation in photoprotective traits. For each model/year combination, MTA analyses were conducted using the sorghum SNP set (GWAS) and transcript expression data (TWAS). Genes in LD with the top 0.05% of GWAS SNPs (Supporting information Table S3) and the top 1% of TWAS genes (Supporting information Table S4), as ranked by Bonferroni-adjusted *p*-value, were brought forward as “top” genes. Additionally, genes in LD with the top 10% of GWAS SNPs and their corresponding transcript expression levels were analysed for covariance with the NPQ traits via Fisher’s Combined Test (FCT), with the top 1% of genes by Bonferroni-adjusted *p*-value brought forward as top FCT results (Supporting information Table S5).

An intersection-based approach was used to produce a list of candidate genes that control sorghum photoprotection traits. As an example for a single trait, maximum NPQ may be reasonably representative of an accession’s photoprotective capacity, particularly as it tended to correlate significantly with both NPQ induction and relaxation *k*. The joint model GWAS for maximum NPQ identified four SNPs with an FDR-adjusted *p*-value below 0.05, spanning a region of less than 20 kilobases within two LD blocks containing 74 genes (Figure 8 **a)**. When combined with TWAS results (Figure 8 **b & c**) for the joint analysis, the FCT tests in both GP and 3L tissues (Figure 8 **d & e**) strengthened this association with several genes in the same locus comprising some of the highest-confidence genes in the analysis. This region was also highly enriched in the joint FCT of CT1, the multivariate combination of maximum NPQ, NPQ induction amplitude, and NPQ induction and relaxation *k*. A number of top genes overlapped in multiple joint-model maximum NPQ analyses (Figure 8 **f**), with five genes overlapping in three separate analyses and 93 genes overlapping in two separate analyses. Seventy genes, overlapping in eight or more model analyses for individual traits, were considered progressively higher confidence based on total number of overlaps. The 16 genes overlapping in nine or more analyses are summarised in Table 2.

**Figure 8:**
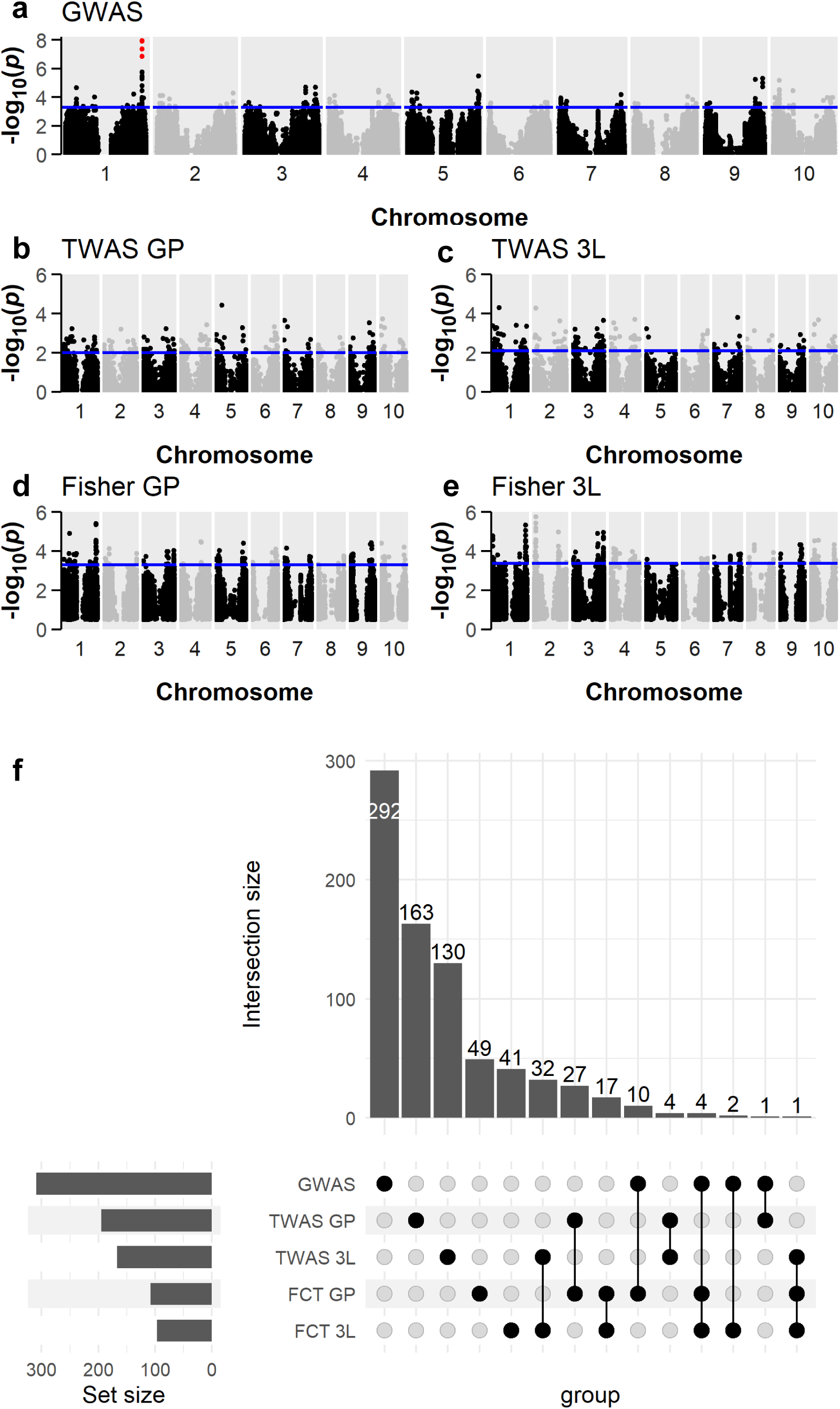
Chromosome mapping (physical location) for SNPs and genes associated with maximum NPQ combined 2017/2019 adjusted means (a-e) and upset plot (f) showing the number of overlapping genes between the top 1% of hits in FCT 3L and GP, TWAS 3L and GP, and top 0.05% of hits in GWAS analysis, for maximum NPQ (joint model). a, GWAS; b, TWAS in GP tissue; c, TWAS in 3L tissue; d, Fisher’s combined test from GP tissue; e, Fisher’s combined test from 3L tissue; Blue lines indicate the threshold of SNPs in top 0.05% (a) or genes in top 1% (a-e) of -log_10_ *p* values. SNPs in plot A with an FDR-adjusted *p*-value <0.05 are highlighted in red. TWAS and FCT gene positions are plotted as midpoint of each gene.

**Table 2:**
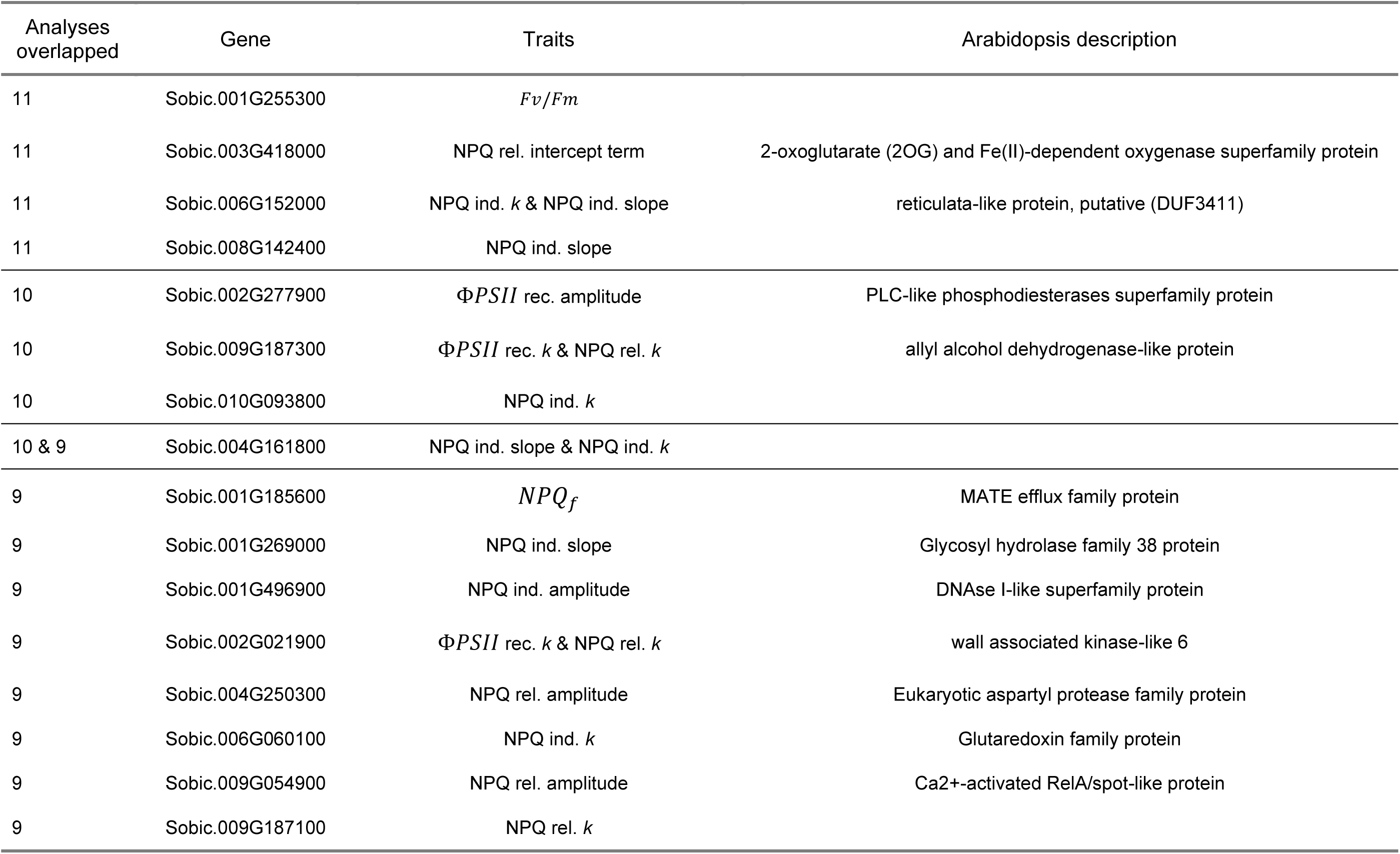
Top sorghum genes overlapping in nine or more analyses for various NPQ kinetic traits.

Many top genes were common to multiple traits, with nearly 30% of top genes overlapping three or more trait top gene lists. The 37 genes which overlapped 10 or more traits were considered candidates; Table 3 summarises the highest-confidence portion of those. The multi-model/single trait and multi-trait overlap thresholds together resulted in 104 unique candidates (Supporting information Table S2). An additional six unique genes were recorded as candidates based on overlapping associations with three or more traits and being either annotated in TAIR for light-response related function or noted as a carotenoid biosynthesis prior in Ortiz et al. (2017).

**Table 3:**
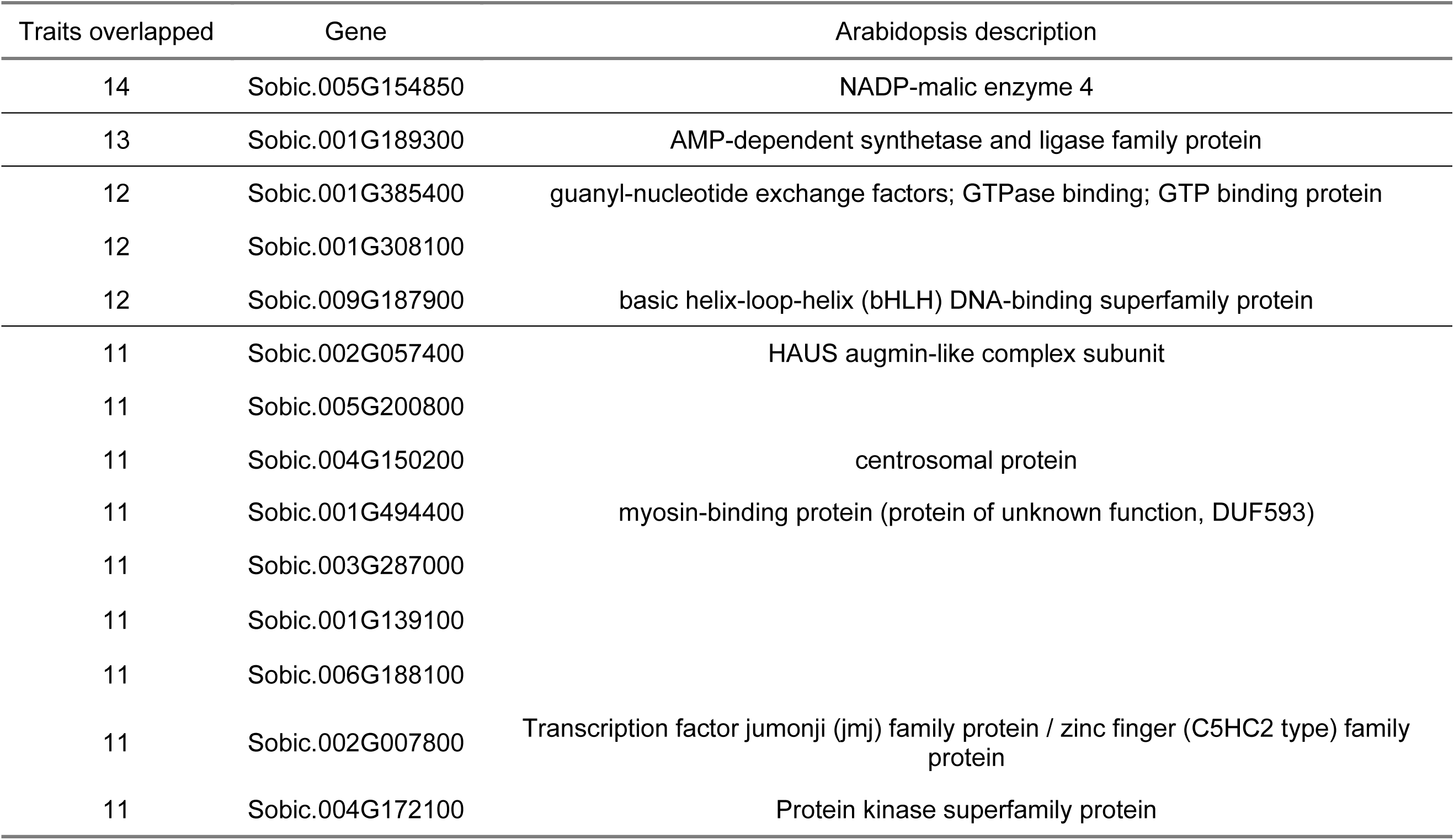
Genes in the top 0.05% of GWAS and top 1% of TWAS and FCT analyses overlapping 11 or more traits.

Several of the 104 candidates are orthologous to Arabidopsis genes which based on their annotation could contribute directly or indirectly to photosynthetic efficiency and photoprotective traits in dynamic light conditions. Sobic.006G152000 overlapped in 11 separate analyses for NPQ induction slope and *k*, eight analyses for CT3 (max NPQ, NPQ induction *k*, and NPQ relaxation *k*) and several analyses for the other combined traits. The Arabidopsis orthologue (*LCD1*) of this gene is involved in palisade mesophyll cell density (Barth & Conklin, 2003). Sobic.003G418000, which overlapped 11 analyses for the NPQ relaxation intercept term, has been suggested to alter strigolactone biosynthesis in response to parasite stress in sorghum (Bellis et al., 2020), and is orthologous to Arabidopsis LBO1, involved in strigolactone biosynthesis (Brewer et al., 2016). Similarly, Sobic.006G060100 (overlaps nine analyses for NPQ induction *k*) is orthologous to Arabidopsis AT5G58530, a glutaredoxin (GRX)-like protein.

Sobic.009G187300, with 10 overlaps for NPQ relaxation rate constant and the correlated 𝛷𝑃𝑆𝐼𝐼 recovery rate constant, is an orthologue of Arabidopsis AT3G59840, an allyl alcohol dehydrogenase-like protein which has been implicated in regulation of ATP synthase activation/deactivation kinetics (Gong et al., 2006). Sobic.001G185600, overlapping nine analyses for 𝑁𝑃𝑄_𝑓_, is orthologues to a MATE efflux family protein in Arabidopsis (AT1G11670) which has been associated with light-induced regulation of gene expression during etiolation, potentially playing an early role in preparing Arabidopsis seedlings for photosynthesis (Hudson et al., 2003).

Of the multiple-trait overlap candidates, Sobic.005G154850 was a top gene in 14 traits analysed, predicted as an NADP-ME isoform (not apparently involved in the C4 carbon concentrating mechanism) and orthologous to AT1G79750 (NADP-ME4). Sobic.001G189300, orthologous to an AMP-dependent synthase and ligase family protein, overlaps 13 top gene sets. This jasmonic acid precursor has been implicated in stress response in Arabidopsis (Bonsegna et al., 2005). Notably, Sobic.006G128300 overlapped as a top gene in seven separate analyses and is orthologous with the Arabidopsis *NPQ6* gene, a YCF-20 family gene involved in energy-dependent quenching (Jung & Niyogi, 2010). Additionally, Sobic.008G021000 was found to overlap four top gene analyses for both NPQ relaxation *k* and 𝛷𝑃𝑆𝐼𝐼 recovery *k*. This gene was also identified in a genetic screen for fluorescence parameters in sorghum (Ortiz et al., 2017) and annotated as *PHYTOCHROME INTERACTING FACTOR 4* (PIF4), a transcriptional regulator of carotenoid biosynthesis (Ruiz-Sola et al., 2014).

SIFT analyses of coding sequence SNPs within candidate genes were performed to uncover potential causal SNPs which may play a part in photoprotective regulation (Supporting information Table S8). Of the 110 candidate genes, 99 contained at least one predicted nonsynonymous substitution (encoding a different amino acid than the reference allele), providing one possible causal mechanism underlying the observed marker-trait associations.

## Discussion

Fine-tuning photoprotection to better match prevailing light conditions could contribute toward improvement in photosynthetic efficiency in the crops that feed and fuel the world (Zhu et al., 2010). Here, variation in NPQ in a large diversity panel was screened during two field seasons, confirming significant heritable variation in NPQ resides within the sorghum genome. Subsequent genome- and transcriptome-wide analyses identified a number of candidate loci underlying genetic control of NPQ in one of the world’s most stress-resilient crops.

### The sorghum genome exhibits wide-ranging and heritable capacity for NPQ

The variation in photoprotective traits observed within this sorghum population adds to a growing body of evidence that significant intraspecies variability in photoprotective capacity exists in C4 crops. BLUPs of maximum NPQ measured during the 10-minute high light treatment varied from just above 2.4 up to more than 3.2, a somewhat smaller range than that observed in field-grown sorghum and maize panels subjected to similar light treatment conditions (Ferguson et al., 2025; Sahay et al., 2023, 2024) but similar to that found in a panel of 529 diverse rice accessions investigated by Wang et al. (2017), who reported mean NPQ values of 2.4 to 3.0 measured in the field after imposition of five minutes of 1,000 μ𝑚𝑜𝑙 m^-2^ s^-1^ PPFD. Additional studies in rice diversity panels also noted genetic variation in NPQ expressed either as a quantum yield (Quero et al., 2021) or as NPQt (Wei et al., 2022), a simplified NPQ parameter that does not require dark-adaptation. Herritt et al. (2016) reported differences in NPQ between accessions of a diverse soybean panel, derived via measurements of photochemical reflectance index, which is correlated with the energy-dependent *qE* component of NPQ. NPQ determined from high-throughput fluorescence imaging of Arabidopsis rosettes ranged from 1.5 to 4 (non-averaged values) in contrasting accessions after eight minutes at 1,000 μ𝑚𝑜𝑙 m^-2^ s^-1^ PPFD (Rungrat et al., 2019). This is similar to the quality-filtered unaveraged sorghum NPQ values in the current study which ranged from 1.0 to 4.3. While actinic light intensities and treatment timing differ between these studies, it is clear that variation in NPQ exists within single species, including economically important crops.

NPQ, like many photosynthetic traits, is developmentally and environmentally plastic (Bielczynski et al., 2017). Accordingly, the distribution of several trait values in this study shifted between 2017 and 2019 (Figure 4), though notably maximum NPQ was more correlated between both years than most of the other NPQ parameters. The faster NPQ induction and slower NPQ relaxation rates, and higher residual 𝑁𝑃𝑄_𝑓_in 2017 compared to 2019 suggest a “more protected” state in 2017. The differences between years may be explained in part by less frequent and lower total precipitation during the early part of the 2017 growing season (Supporting information Figure S1). In a similarly diverse multi-year sorghum NPQ screen, Sahay et al. (2024) observed stronger correlations in NPQ kinetics between years than between low and high nitrogen treatments imposed within years, providing further evidence of the environmental plasticity of NPQ in the field. In the current study, the resulting reduction in water availability may have reduced downstream photosynthetic capacity, leading to upregulation of NPQ responses. Indeed, the depressed 𝑃𝐼 and *Fv*/*Fm* values measured in 2017 suggest a higher level of photoinhibition, despite the increased induction of sustained NPQ. Despite this plasticity, the medium-to-high levels of heritability found for max NPQ and induction and relaxation rate constants in this and other studies, and significant correlations between different years, facilitated the use of GWAS and TWAS to determine underlying causal genes.

The strong positive correlation observed here between BLUPs for NPQ relaxation *k* and 𝛷PSII recovery *k* indicates that these two traits may be genetically and mechanistically linked; similar conclusions based on overlapping candidate genes for NPQ and PSII traits, associated with control of photosynthetic efficiency, support this linkage (Sahay et al., 2023). Faster NPQ relaxation rate constants were positively correlated with lower photoinhibition during the light treatment, as evident from the positive, significant correlation between 𝑃𝐼 and NPQ relaxation *k* in 2017. Arabidopsis mutants with faster NPQ relaxation kinetics during a similar length of treatment showed similar results (X.-P. Li et al., 2002), though the particularly high maximum NPQ reached by PsbS overexpression mutants in that study may also have played a role in photoinhibition avoidance. In the current study, maximum NPQ reached was not correlated with 𝑃𝐼, suggesting that during this short-term high light treatment, NPQ induction kinetics were a larger determinant of potential photoinhibition than the actual level of NPQ attainable by a given accession.

### Genes related to stress response and photosynthetic control are newly associated with NPQ traits

To increase confidence in identifying loci with genes that likely play a role in controlling photoprotection in sorghum, we employed an ensemble approach combining GWAS, TWAS, and FCT analyses as well as leveraging the covariance between fluorescence parameters. Genes which overlapped in eight or more analyses for a given trait, or were top genes for 11 or more traits, were considered candidate genes. Several of the candidates are orthologous to Arabidopsis genes annotated for light use or photosynthesis-related processes. The Arabidopsis *LCD1* gene, similar to Sobic.006G152000, is involved in palisade mesophyll cell density, and mutant phenotypes exhibit sensitivity to growth light conditions, though without an apparent response to short-term high-light stress (Barth & Conklin, 2003). If the gene function is conserved in sorghum, modulation of cell density may affect light absorption and photosynthetic capacity, both of which could impact NPQ. Sobic.003G418000 is involved in strigolactone biosynthesis (Bellis et al., 2020). Strigolactones regulate plant growth responses to suboptimal conditions and share a biosynthesis pathway with carotenoids involved in photoprotection and antioxidant defence (Hirschberg, 2001). Recently, Thula et al. (2022) also observed a direct role of strigolactones in influencing high light tolerance of photosynthesis in Arabidopsis. Sobic.006G060100 is orthologous to AT5G58530 in Arabidopsis, a GRX-like protein which could be involved in abiotic stress responses via redox regulation and antioxidant capacity (Rouhier et al., 2008). Sahay et al. (2023) also uncovered multiple thioredoxin candidate genes associated with NPQ in maize. It is not surprising to find that redox metabolism and antioxidant scavenging are deeply intertwined with both short and longer-term photoprotective processes (Foyer et al., 2012; Müller-Moulé et al., 2002), with overlap in mechanisms as well as precursor molecules. Variability in dynamic photoprotective responses due to underlying variation in antioxidant scavenging may therefore be manifested here.

Interestingly, Sobic.008G142400, previously identified as a triterpenoid synthesis gene involved in cuticular wax formation (Busta et al., 2021), was a strong candidate gene in this study overlapping 11 separate analyses. While Sahay et al. (2024) observed only seven of the same candidate genes in their sorghum NPQ screen as found here, notably, Sobic.008G142400 was one of the common candidates. Correlations between cuticular wax and NPQ (and other photosynthetic traits) have been previously noted in other species but with rather inconsistent results (W. Li et al., 2023; Vijayaraghavareddy et al., 2022). This suggests that a mechanistic study between cuticular wax and photoprotection may be warranted to specifically investigate the degree to which photoprotective capacity is altered by cuticular wax coverage, or vice versa. As wax amount and composition have been shown to be heritable traits (Haque et al., 1992; Tassone et al., 2016), optimization of these traits for specific crop environments might provide another potential route toward improving photoprotective and photosynthetic efficiency.

Sobic.009G187300 is orthologous to AT3G59840, an allyl alcohol dehydrogenase-like protein, which plays a role in activation and deactivation of ATP synthase (Gong et al., 2006). Variation in ATP synthase activation kinetics may have affected NPQ via effects on the chloroplast stroma/thylakoid lumen pH gradient, given the role of lumenal pH on PsbS and xanthophyll cycle-dependent NPQ (X.-P. Li et al., 2000). Sobic.001G269000, orthologue of Arabidopsis AT5G66150, overlapped nine analyses for NPQ induction slope. The gene is linked to N-glycosylation modification in Arabidopsis (Strasser et al., 2006); though N-glycosylation ubiquitously affects protein biogenesis and function in plants (Strasser, 2022), mutants in N-glycosylation have shown impairment in NPQ capacity and quantum efficiency (Jiao et al., 2020).

Of the genes overlapping a high number of traits, Sobic.005G154850 is an interesting candidate, orthologous to several NADP-ME isoforms in Arabidopsis which have been implicated in response to diverse stressors and in developmental plasticity (Badia et al., 2015; Voll et al., 2012). One such orthologue is the fatty-acid oxidation pathway NADP-ME4 associated with photomorphogenesis in the ancestral C_3_ (Ma et al., 2002); variability in this enzyme could conceivably be responsible for developmental plasticity in light harvesting and photoprotective capacity. This locus was recently found in a diverse sorghum panel to be associated with growth rate and plant height (Panelo et al., 2024). Sobic.005G154850 is a truncated protein with a large mis-sense section at the 3’ end, but it is unclear if this could have affected NADP-ME activity directly involved in the C_4_ decarboxylation cycle. Sobic.001G189300 is orthologous to a jasmonic acid precursor and overlapped 13 top gene lists. Jasmonic acid (JA) and its precursors have a well-defined role in antioxidant stress response (Munné-Bosch, 2005; Yamauchi & Matsushita, 1979); within-population variance in JA production and activity could quite reasonably underlie variability in the photoprotection traits measured here. Nearly 40 candidate genes spread across several chromosomes overlapped ten or more individual NPQ and PSII kinetic traits, suggesting that variation in single loci can functionally affect multiple aspects of photoprotection and photochemistry, a pattern also observed in Sahay et al. (2024).

The previously discussed genes are mostly novel candidate genes associated with variation in sorghum photosynthetic efficiency in dynamic light conditions. While large-scale quantitative genomic studies like this are not causal evidence for functional regulation, they shed light on genomic regions that have strong associations with the traits of interest, providing a valuable resource for future validation studies. In addition to the candidates listed above, several candidate genes were found to be orthologous to Arabidopsis genes previously associated with photoprotection. An orthologue to Arabidopsis chloroplast lipocalin LCNP (Sobic.006G222700) was identified in the overlap between 6 different analyses for three separate traits including NPQ relaxation slope and 𝑁𝑃𝑄_𝑓_. The gene product from the Arabidopsis ortholog is involved in sustained NPQ (qH, Malnoë et al., 2017), and its association with residual NPQ after light treatment in this study suggests it may function similarly in sorghum. Sobic.006G128300 overlapped seven trait top gene lists and is orthologous to AT5G43050, a chloroplast-localised YCF20 gene sharing 70% sequence similarity (Berardini et al., 2015). SIFT analysis found two nonsynonymous SNPs in the coding sequence of this gene in Sorghum, which is especially interesting since an Arabidopsis mutant carrying a single base-pair deletion within this gene (mutant *NPQ6*) exhibits reduced NPQ (Jung & Niyogi, 2010). SIFT analysis across the 110 candidate genes showed that SNPs within coding regions of 90% of the identified candidates were predicted to contain nonsynonymous substitutions, which provides a potential mechanism which could give rise to the observed phenotypic variation. A similarly high prevalence of nonsynonymous substitutions was found across all genes in a comparative study of deleterious mutations in sorghum and maize (Lozano et al., 2021); the high percentage of nonsynonymous substitutions in photoprotection-associated genes may provide a potential mechanism for some of the heritable diversity in photoprotection within this diverse sorghum population. Given the wide-ranging climatic origins of the panel (Ferguson et al., 2021), nonsynonymous coding region variation in candidate gene sequences may reflect environmental adaptation of NPQ characteristics. The complex and diverse genetic architecture underlying dynamic photoprotective traits, along with the measured trait heritability, suggests that informed genomic selection could affect meaningful changes in NPQ phenotypes in sorghum.

## Conclusion

This work characterised the extent of heritable variation in photoprotective traits in sorghum using the largest field-grown diversity panel for these parameters to date. The results highlight the complexity of genetic control of photoprotection in this species, with a wide range of significant small effect loci associated with the fluorescence parameters. Using an ensemble approach combining GWAS and TWAS and leveraging the covariance between fluorescence parameters, several novel high-confidence candidates were identified which may exert genetic control of photosynthesis and photoprotection. This information can be used to inform further, targeted experiments via mutant or genetic modification studies to manipulate NPQ in sorghum, and potentially if combined with appropriate marker development could be used in marker or genomics-assisted breeding for photosynthetic efficiency in sorghum and related C4 crops. Most of the identified candidate genes were not previously annotated with direct functions in photoprotection, but several of the candidates are associated with adjacent processes that are indirectly linked to photosynthesis and photoprotection. While germplasm-focused enhancements in crop productivity will require a whole-plant focus (Lopes et al., 2011), better understanding genetic control over photoprotective capacity in challenging stress conditions could play a role in development of crop germplasm better-suited to anticipated hotter, drier conditions across many of the global crop growing regions.

## Acknowledgements

We thank Pat J. Brown at the University of California, Davis for planting the 2017 sorghum plots, and Julian Hibberd and Conor Simpson at the University of Cambridge for helpful comments and discussion on genomic data analysis. This work was supported by funding from the Frank Smart Studentship in Botany to RLV and Gatsby Foundation startup funding to JK.

## Author contributions

RV: Conceptualisation; project administration; investigation; methodology; visualisation; formal analysis; supervision; writing – original draft (lead); writing – review and editing (lead); SF: Investigation; methodology; BM: Investigation; methodology; KG: Investigation; methodology; JW; Investigation, writing – review and editing; AL: Methodology; JF: Conceptualisation; CB: Resources; TP: Resources; JK: Conceptualisation (lead); project administration (lead); investigation; methodology; writing - review and editing; supervision (lead); funding acquisition; resources.

## Data availability

The NPQ trait data that support the findings of this study are openly available in the supporting materials.

## A Supporting information

**SI Figure S1**: Plots of maximum daily temperature and total daily precipitation recorded at the Willard Airport weather station (Savoy, IL, USA) during the 2017 and 2019 sorghum panel growing seasons.

**SI Figures S2-S5**: Chromosome mapping (Manhattan) plots for single nucleotide polymorphisms associated with NPQ and combined NPQ traits in 2017, 2019, and joint genome-wide association study analyses.

**SI Figures S6-S11**: Chromosome mapping (Manhattan) plots for genes associated with NPQ and combined NPQ traits in 2017, 2019, and joint transcriptome-wide association study analyses in third-leaf and growing point tissue.

**SI Figures S12-S17**: Chromosome mapping (Manhattan) plots for genes associated with NPQ and combined NPQ traits in 2017, 2019, and joint Fisher’s combined test analyses in third-leaf and growing point tissue.

**SI Table S1**: Linkage disequilibrium (LD) blocks containing top 0.05% of SNPs from GWAS.

**SI Table S3**: Summary of candidate genes

**SI Table S4**: Top 1% of genes identified via TWAS for each trait, tissue, and model year.

**SI Table S5**: Top 1% of genes identified via Fisher’s combined test for each trait and model year.

**SI Table S6**: Gent ontology (GO) biological process enrichment analysis of *Arabidopsis thaliana* orthologues of the top sorghum genes which overlapped in three or more analyses

**SI Table S7**: Summary of genes in top results for FCT 3L, FCT GP, TWAS 3L, TWAS GP (top 1%), and GWAS (genes in LD with top 0.05% of SNPs), for each analysis

**SI Table S8**: SIFT scores of coding sequence single nucleotide polymorphisms SNPs within candidate genes.

**SI Table S9**: BLUPs of all traits for each accession and model year.

**SI Table S10**: 2017 NPQ trait values for individual leaf discs

**SI Table S11**: 2019 NPQ trait values for individual leaf discs

