## Supplemental Figures S1-17 for "High throughput screen of NPQ in sorghum shows highly polygenic architecture of photoprotection"

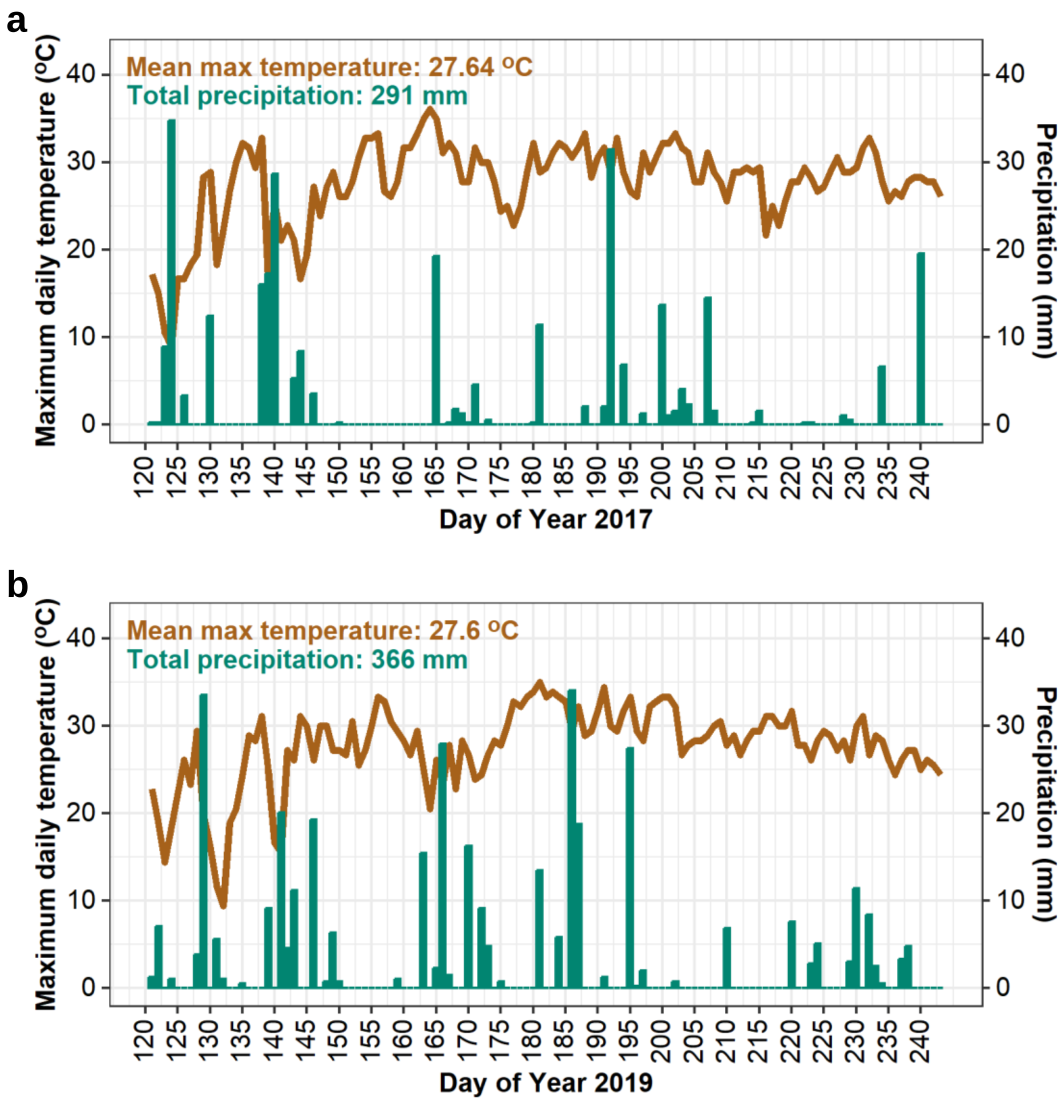


**Figure S1: Plots of maximum daily temperature (brown lines) and total daily precipitation (green columns) recorded at the Willard Airport weather station (Savoy, IL, USA) during the 2017** (a) **and 2019** (b) **sorghum panel growing seasons. Seasonal totals calculated from May 05 through August 31. Planting date: 151. Sampling dates: 206-216 (2017); 203-212 (2019). Data retrieved from www.ncdc.noaa.gov (Station ID GHCND:USW00094870; 3.1 km from Maxwell Farm and 6.73 km from Energy Farm).**

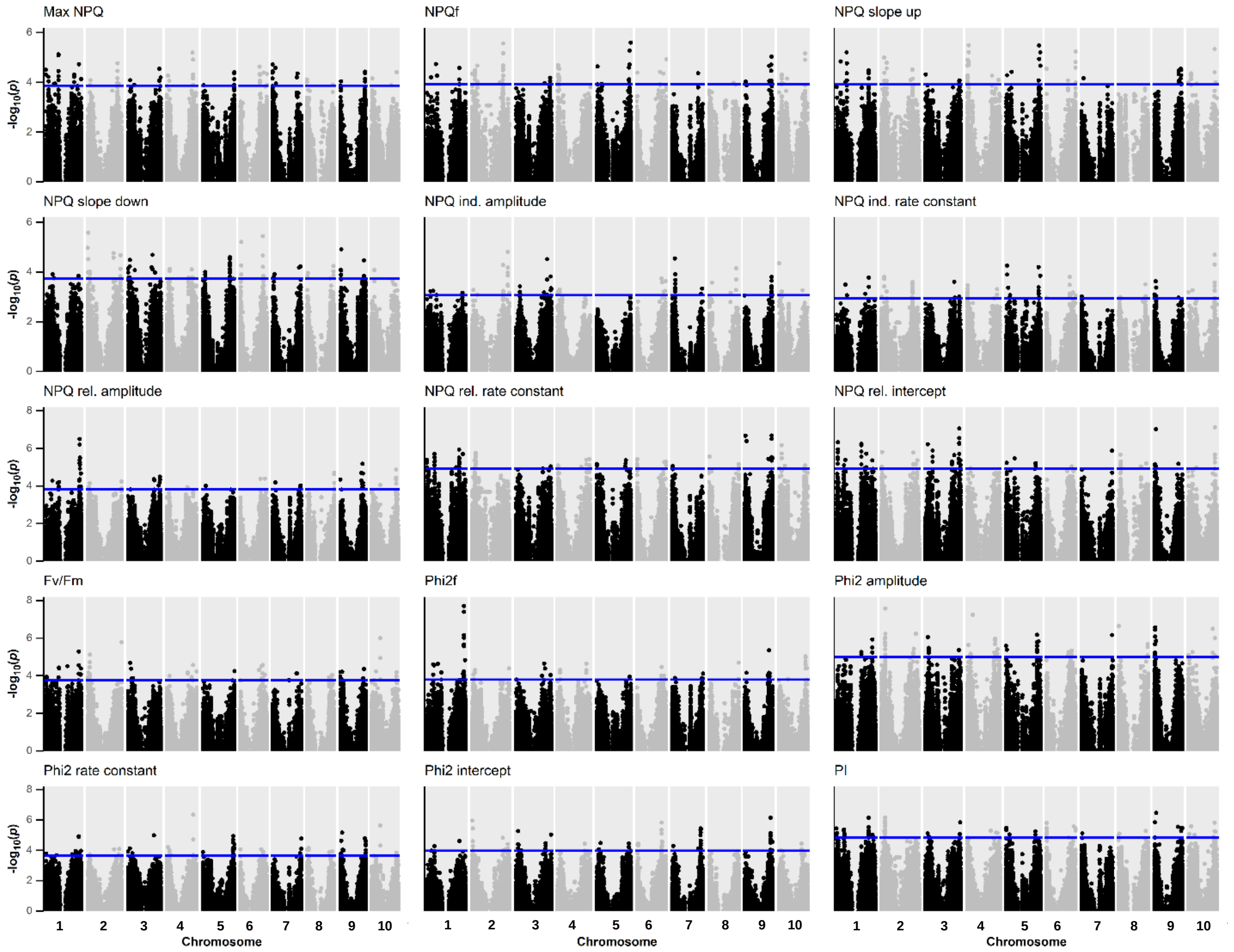


**Figure S2: Chromosome mapping (physical location) for single nucleotide polymorphisms (SNPs) associated with 2017 genome-wide association study (GWAS) non-photochemical quenching traits. Blue line indicates threshold of SNPs in top 0.05% by $-\log_{10}$ *p*-value. 65% of SNPs below *p*-value of 2.5 have been randomly removed from each GWAS plot to reduce image size. NPQf: Final dark time point NPQ value. Phi2: Photosystem II quantum yield. Phi2f: Final dark time point Phi2. PI: Photoprotection index.**

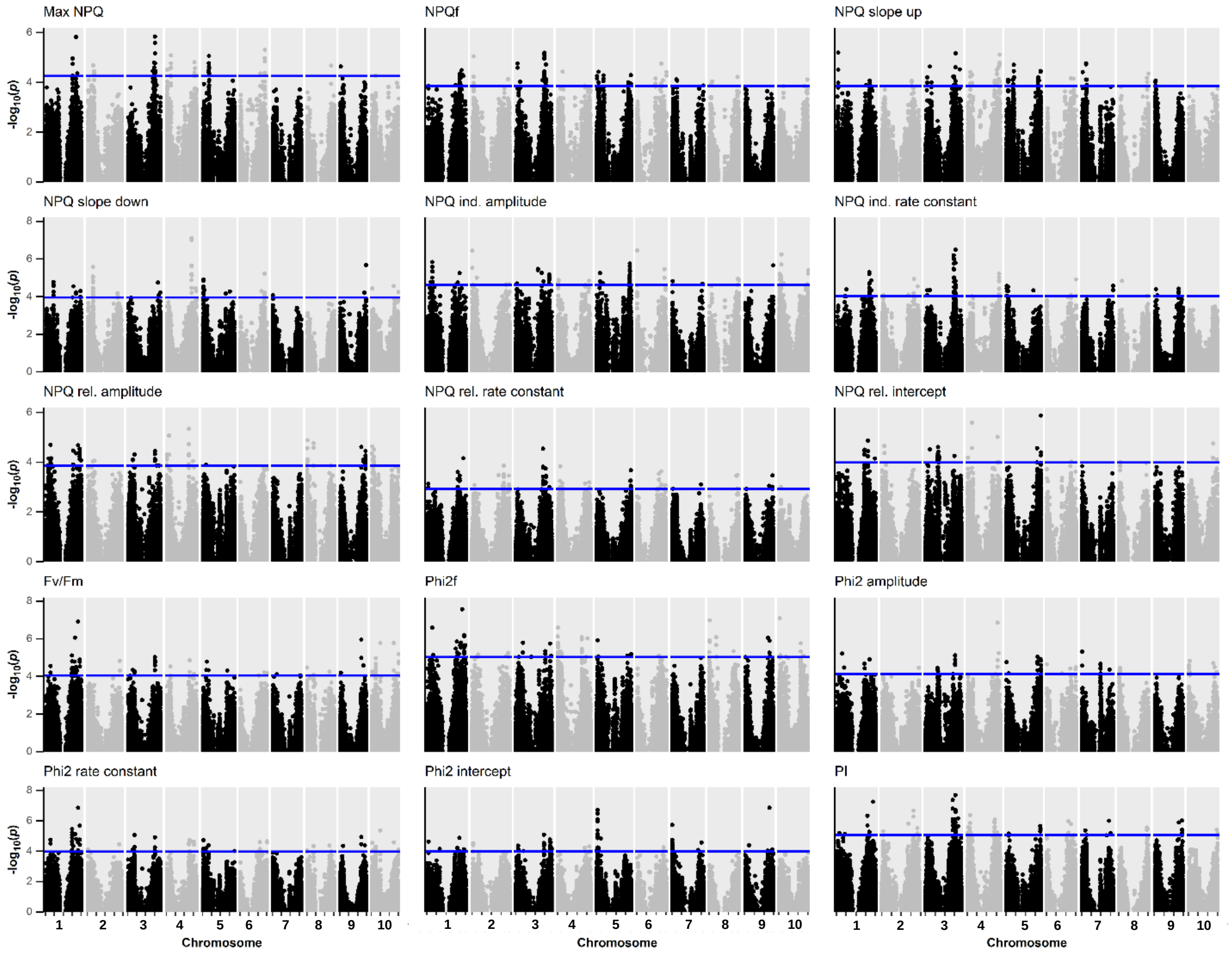


**Figure S3: Chromosome mapping (physical location) for single nucleotide polymorphisms (SNPs) associated with 2019 genome-wide association study (GWAS) non-photochemical quenching traits. Blue line indicates threshold of SNPs in top 0.05% by -$\log_{10}$ *p*-value. 65% of SNPs below *p*-value of 2.5 have been randomly removed from each GWAS plot to reduce image size. NPQf: Final dark time point NPQ value. Phi2: Photosystem II quantum yield. Phi2f: Final dark time point Phi2. PI: Photoprotection index.**

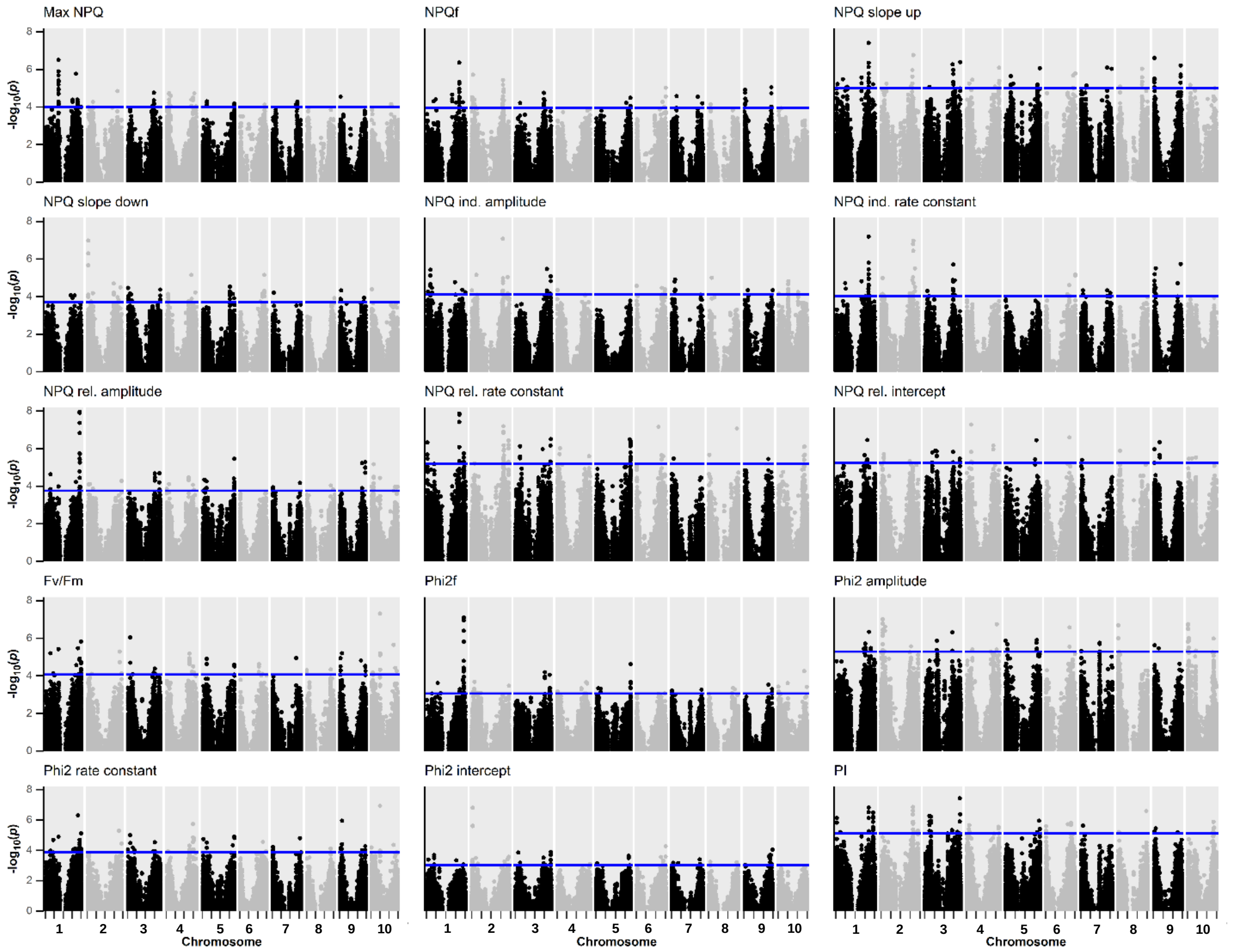


**Figure S4: Chromosome mapping (physical location) for single nucleotide polymorphisms (SNPs) associated with joint genome-wide association study (GWAS) non-photochemical quenching traits. Blue line indicates threshold of SNPs in top 0.05% by -$\log_{10}$ *p*-value. 65% of SNPs below *p*-value of 2.5 have been randomly removed from each GWAS plot to reduce image size. NPQf: Final dark time point NPQ value. Phi2: Photosystem II quantum yield. Phi2f: Final dark time point Phi2. PI: Photoprotection index.**

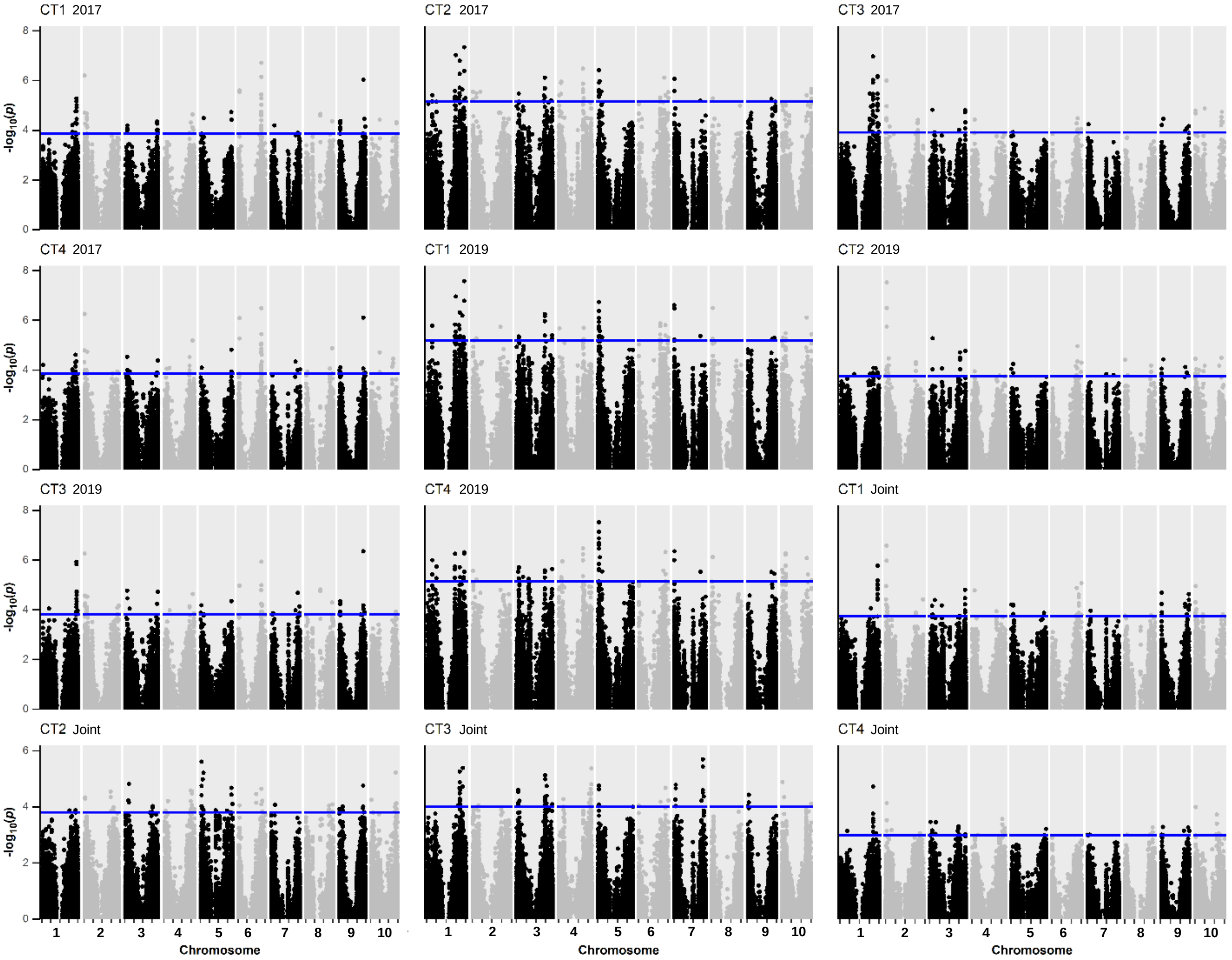


**Figure S5: Chromosome mapping (physical location) for single nucleotide polymorphisms (SNPs) associated with combined-trait (CT) genome-wide association study (GWAS) non-photochemical quenching traits. Blue line indicates threshold of SNPs in top 0.05% by $-\log_{10}$ *p*-value. 65% of SNPs below *p*-value of 2.5 have been randomly removed from each GWAS plot to reduce image size. CT1: Max NPQ, NPQ induction amplitude, NPQ induction rate constant *k*, NPQ relaxation rate constant *k*. CT2: NPQ induction amplitude, NPQ induction *k*, NPQ relaxation rate constant *k*. CT3: Max NPQ, NPQ induction rate constant *k*, NPQ relaxation rate constant *k*. CT4: $PI$, $\Phi PSII$ recovery amplitude, $\Phi PSII$ recovery rate constant *k*.**

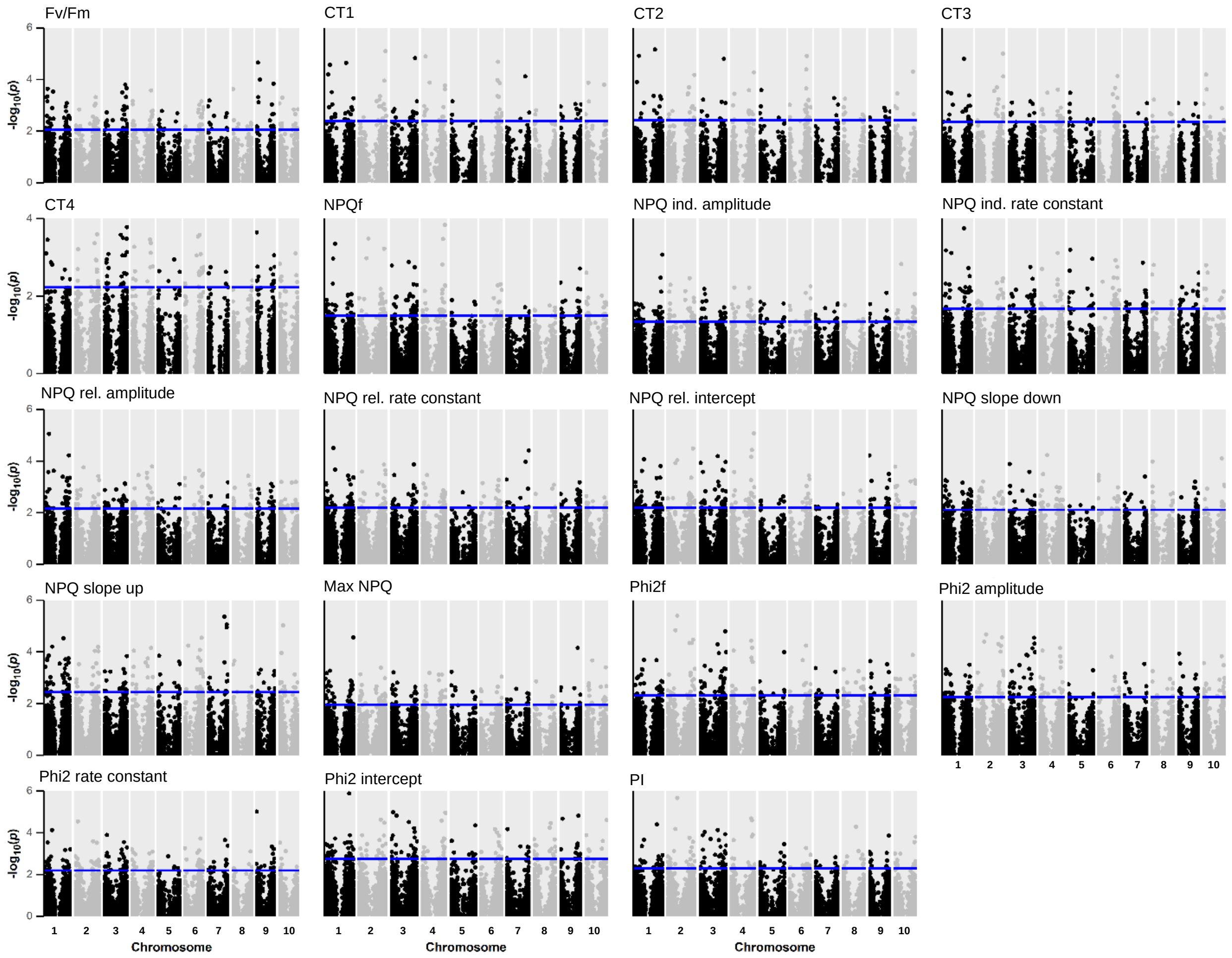


**Figure S6: Chromosome mapping (physical location) for genes associated with 2017 third-leaf tissue transcriptome-wide association study (TWAS) non-photochemical quenching traits. Gene positions plotted as midpoint of each gene. Blue line indicates threshold of genes in top 1% by $-\log_{10}$ *p*-value. CT: Combined trait analysis. CT1: Max NPQ, NPQ induction amplitude, NPQ induction rate constant *k*, NPQ relaxation rate constant *k*. CT2: NPQ induction amplitude, NPQ induction *k*, NPQ relaxation rate constant *k*. CT3: Max NPQ, NPQ induction rate constant *k*, NPQ relaxation rate constant *k*. CT4: $PI$, $\Phi PSII$ recovery amplitude, $\Phi PSII$ recovery rate constant *k*. NPQf: Final dark time point NPQ value. Phi2: Photosystem II quantum yield. Phi2f: Final dark time point Phi2. PI: Photoprotection index.**

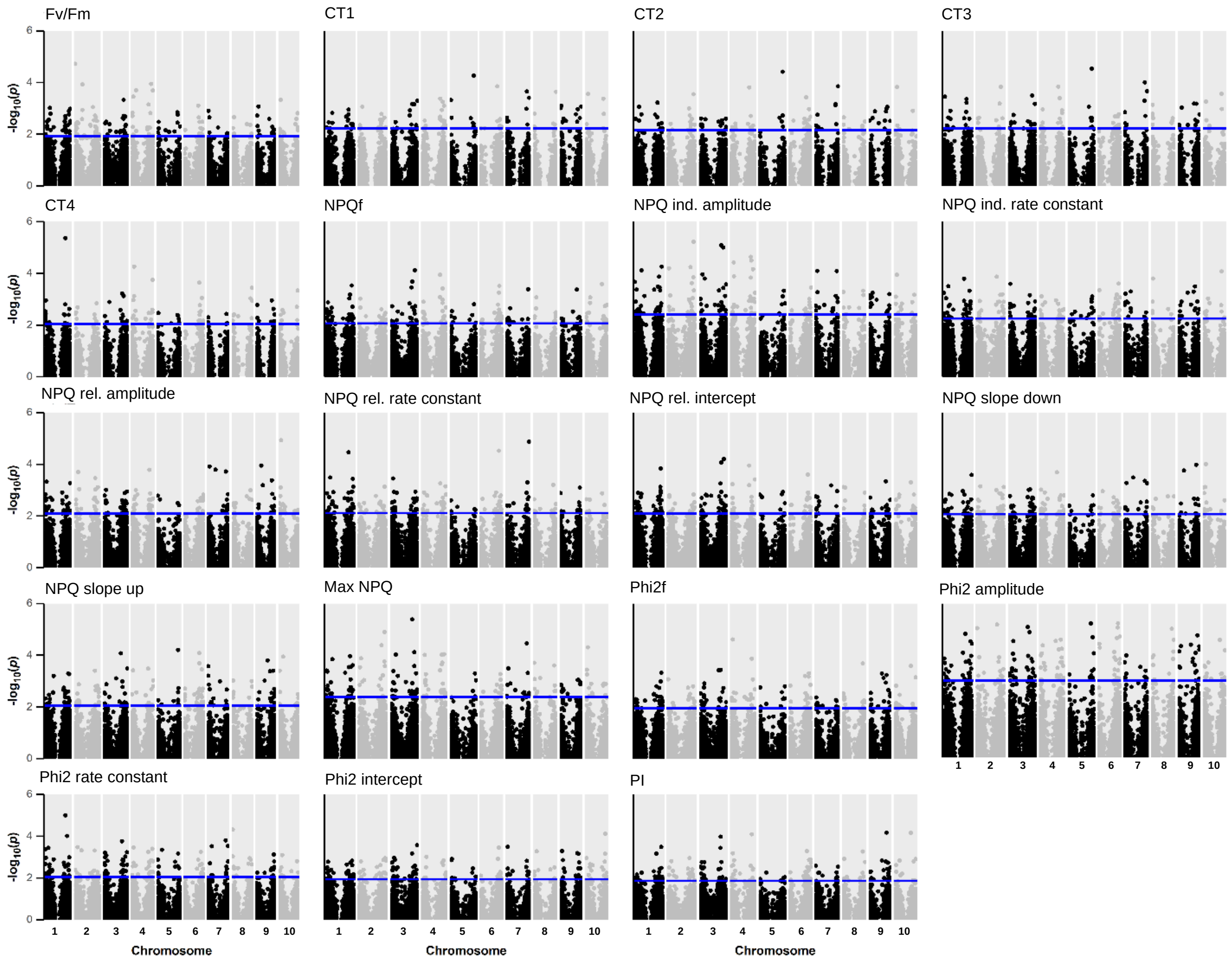


**Figure S7: Chromosome mapping (physical location) for genes associated with 2019 third-leaf tissue transcriptome-wide association study (TWAS) non-photochemical quenching traits. Gene positions plotted as midpoint of each gene. Blue line indicates threshold of genes in top 1% by -$\log_{10}$ *p*-value. CT: Combined trait analysis. CT1: Max NPQ, NPQ induction amplitude, NPQ induction rate constant *k*, NPQ relaxation rate constant *k*. CT2: NPQ induction amplitude, NPQ induction *k*, NPQ relaxation rate constant *k*. CT3: Max NPQ, NPQ induction rate constant *k*, NPQ relaxation rate constant *k*. CT4: *PI*, *ΦPSII* recovery amplitude, *ΦPSII* recovery rate constant *k*. NPQf: Final dark time point NPQ value. Phi2: Photosystem II quantum yield. Phi2f: Final dark time point Phi2. PI: Photoprotection index.**

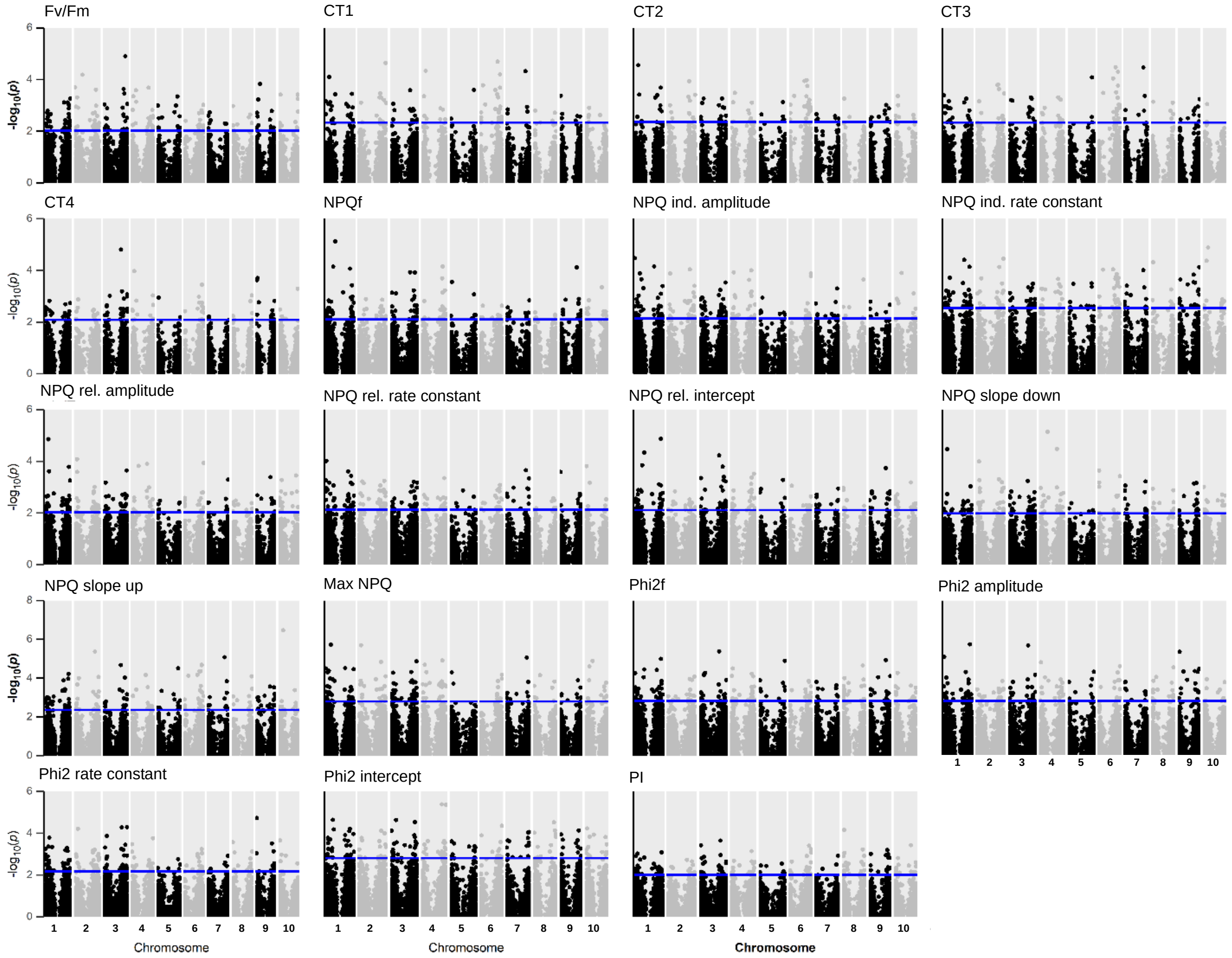


**Figure S8: Chromosome mapping (physical location) for genes associated with joint analysis third-leaf tissue transcriptome-wide association study (TWAS) non-photochemical quenching traits. Gene positions plotted as midpoint of each gene. Blue line indicates threshold of genes in top 1% by -$\log_{10}$ *p*-value. CT: Combined trait analysis. CT1: Max NPQ, NPQ induction amplitude, NPQ induction rate constant *k*, NPQ relaxation rate constant *k*. CT2: NPQ induction amplitude, NPQ induction *k*, NPQ relaxation rate constant *k*. CT3: Max NPQ, NPQ induction rate constant *k*, NPQ relaxation rate constant *k*. CT4: $PI$, $\Phi PSII$ recovery amplitude, $\Phi PSII$ recovery rate constant *k*. NPQf: Final dark time point NPQ value. Phi2: Photosystem II quantum yield. Phi2f: Final dark time point Phi2. PI: Photoprotection index.**

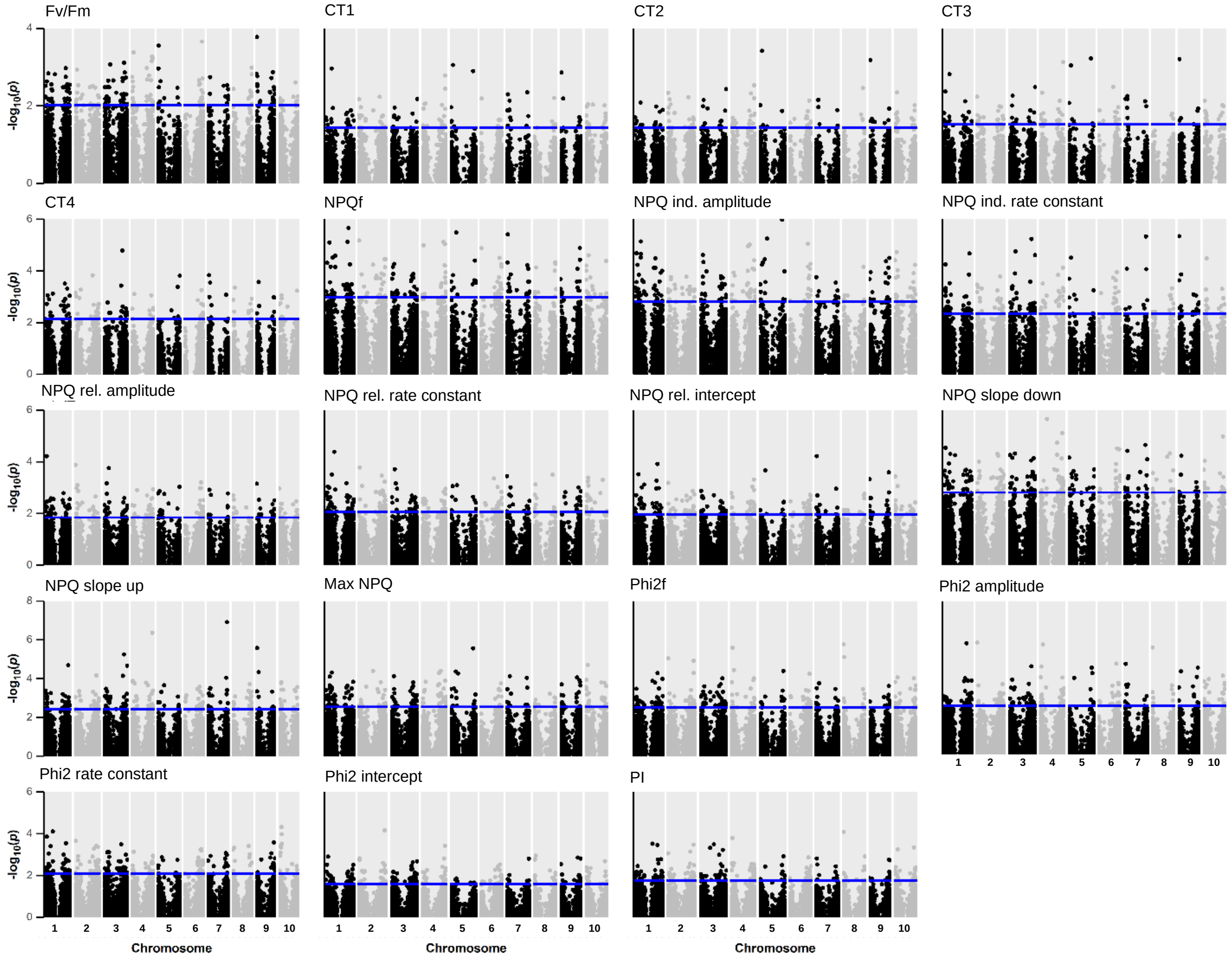


**Figure S9: Chromosome mapping (physical location) for genes associated with 2017 growing point tissue transcriptome-wide association study (TWAS) non-photochemical quenching traits. Gene positions plotted as midpoint of each gene. Blue line indicates threshold of genes in top 1% by -$\log_{10}$ *p*-value. CT: Combined trait analysis. CT1: Max NPQ, NPQ induction amplitude, NPQ induction rate constant *k*, NPQ relaxation rate constant *k*. CT2: NPQ induction amplitude, NPQ induction *k*, NPQ relaxation rate constant *k*. CT3: Max NPQ, NPQ induction rate constant *k*, NPQ relaxation rate constant *k*. CT4: $PI$, $\Phi PSII$ recovery amplitude, $\Phi PSII$ recovery rate constant *k*. NPQf: Final dark time point NPQ value. Phi2: Photosystem II quantum yield. Phi2f: Final dark time point Phi2. PI: Photoprotection index.**

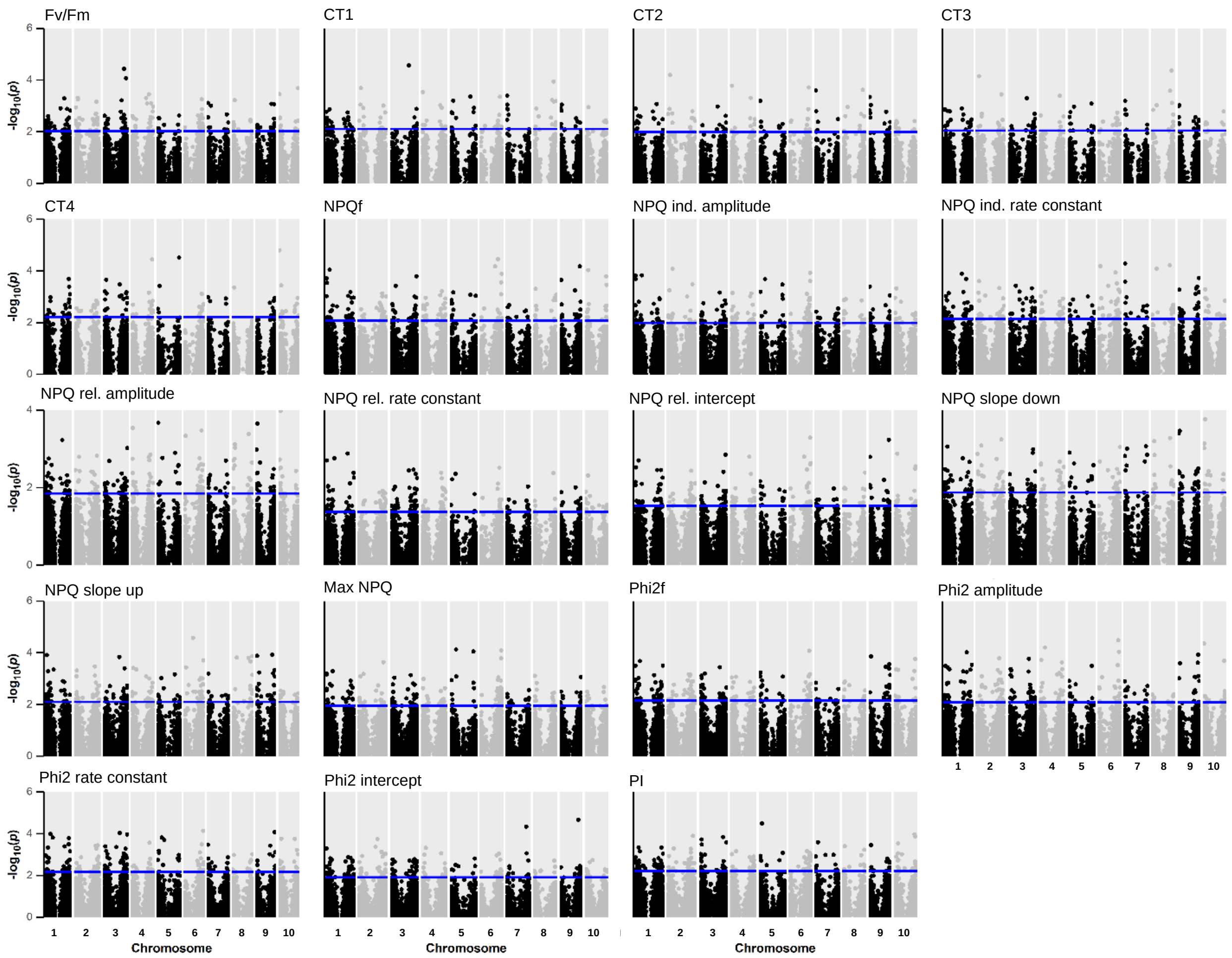


**Figure S10: Chromosome mapping (physical location) for genes associated with 2019 growing point tissue transcriptome-wide association study (TWAS) non-photochemical quenching traits. Gene positions plotted as midpoint of each gene. Blue line indicates threshold of genes in top 1% by $-\log_{10}$ *p*-value. CT: Combined trait analysis. CT1: Max NPQ, NPQ induction amplitude, NPQ induction rate constant *k*, NPQ relaxation rate constant *k*. CT2: NPQ induction amplitude, NPQ induction *k*, NPQ relaxation rate constant *k*. CT3: Max NPQ, NPQ induction rate constant *k*, NPQ relaxation rate constant *k*. CT4: $PI$, $\Phi PSII$ recovery amplitude, $\Phi PSII$ recovery rate constant *k*. NPQf: Final dark time point NPQ value. Phi2: Photosystem II quantum yield. Phi2f: Final dark time point Phi2. PI: Photoprotection index.**

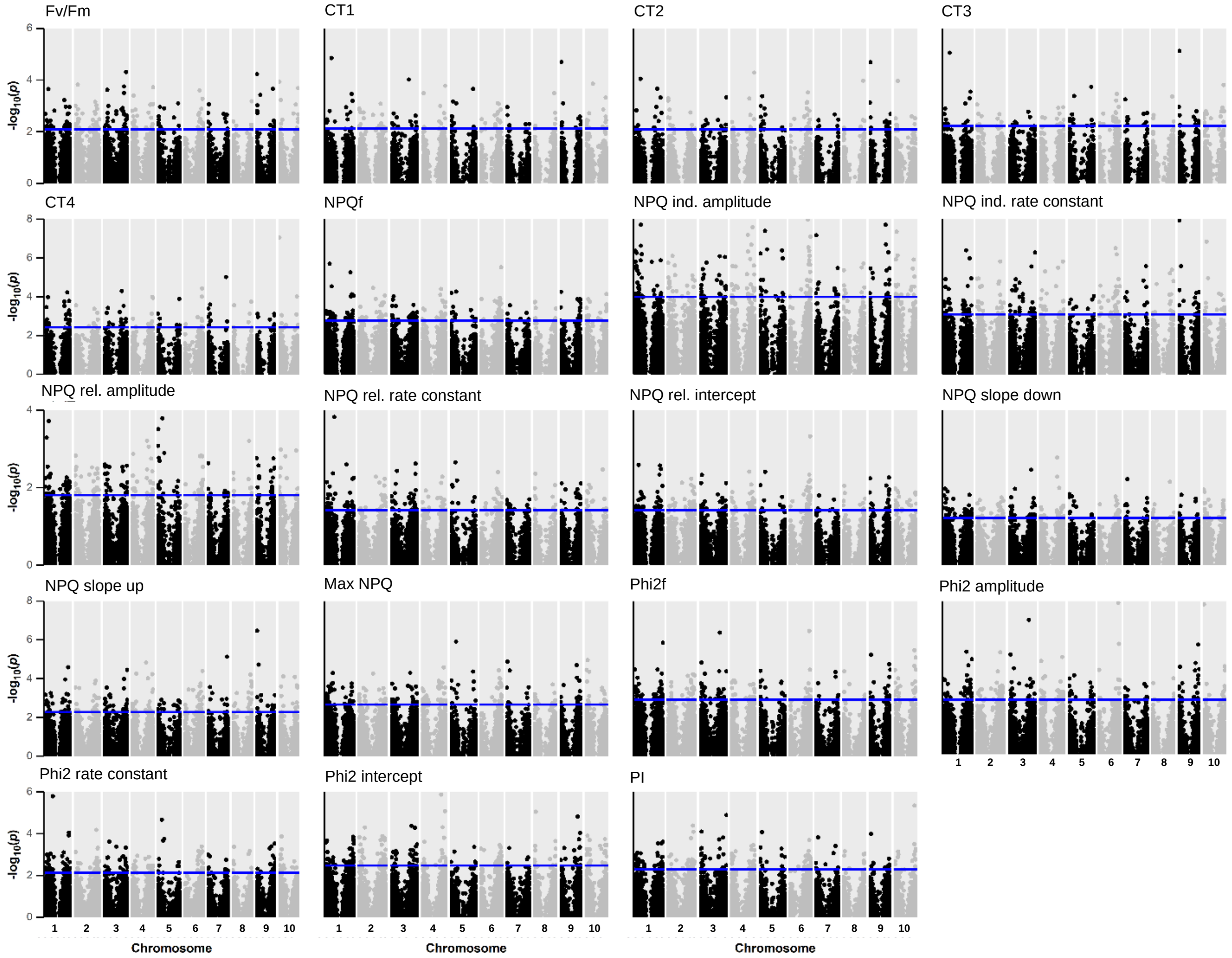


**Figure S11: Chromosome mapping (physical location) for genes associated with joint analysis growing point tissue transcriptome-wide association study (TWAS) non-photochemical quenching traits. Gene positions plotted as midpoint of each gene. Blue line indicates threshold of genes in top 1% by $-\log_{10}$ *p*-value. CT: Combined trait analysis. CT1: Max NPQ, NPQ induction amplitude, NPQ induction rate constant *k*, NPQ relaxation rate constant *k*. CT2: NPQ induction amplitude, NPQ induction *k*, NPQ relaxation rate constant *k*. CT3: Max NPQ, NPQ induction rate constant *k*, NPQ relaxation rate constant *k*. CT4: $PI$, $\Phi PSII$ recovery amplitude, $\Phi PSII$ recovery rate constant *k*. NPQf: Final dark time point NPQ value. Phi2: Photosystem II quantum yield. Phi2f: Final dark time point Phi2. PI: Photoprotection index.**

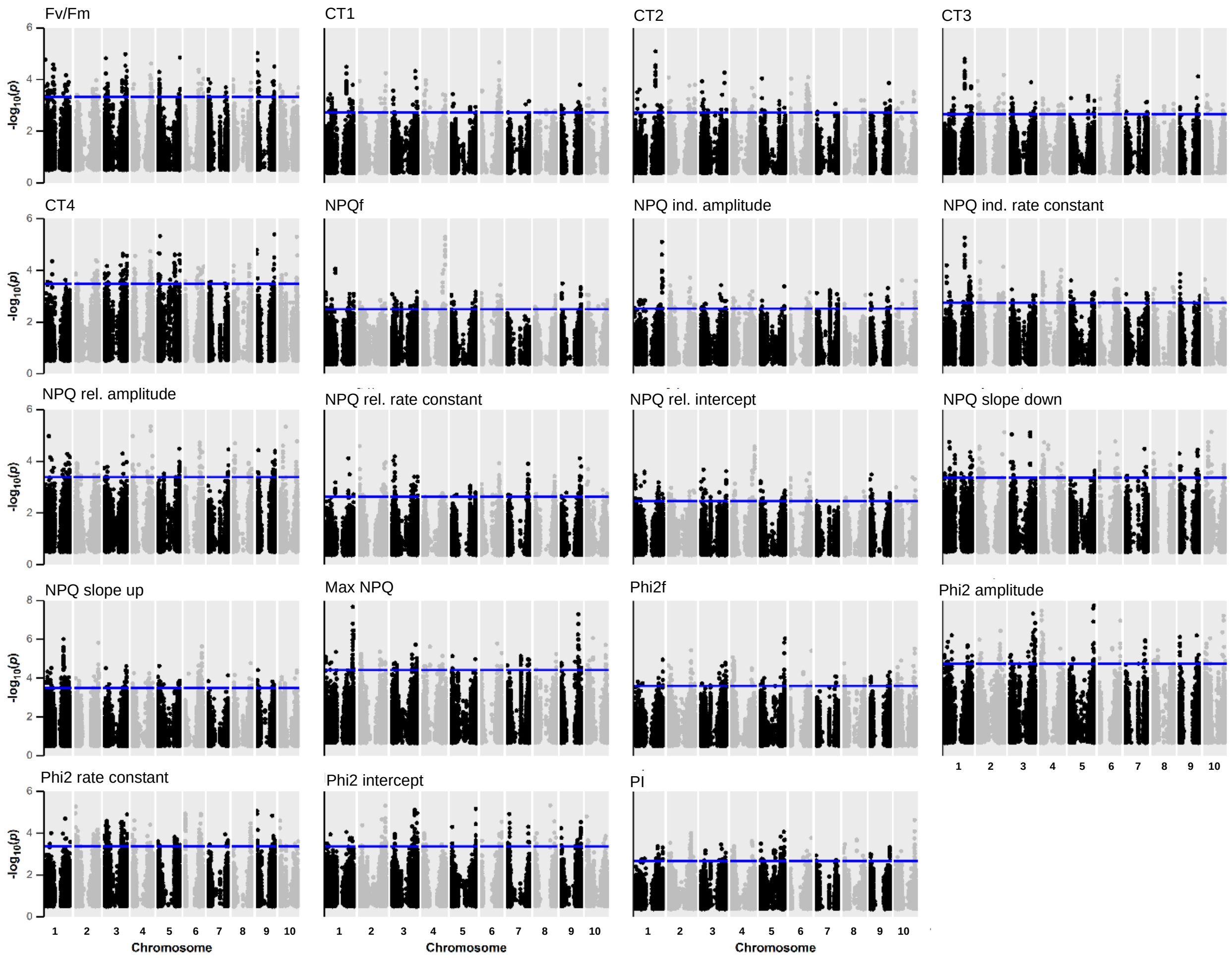


**Figure S12: Chromosome mapping (physical location) for genes associated with 2017 third-leaf tissue Fisher's combined test (FCT) non-photochemical quenching traits. Gene positions plotted as midpoint of each gene. Blue line indicates threshold of genes in top 1% by -$\log_{10}$ *p*-value. CT: Combined trait analysis. CT1: Max NPQ, NPQ induction amplitude, NPQ induction rate constant *k*, NPQ relaxation rate constant *k*. CT2: NPQ induction amplitude, NPQ induction *k*, NPQ relaxation rate constant *k*. CT3: Max NPQ, NPQ induction rate constant *k*, NPQ relaxation rate constant *k*. CT4: $PI$, $\Phi PSII$ recovery amplitude, $\Phi PSII$ recovery rate constant *k*. NPQf: Final dark time point NPQ value. Phi2: Photosystem II quantum yield. Phi2f: Final dark time point Phi2. PI: Photoprotection index.**

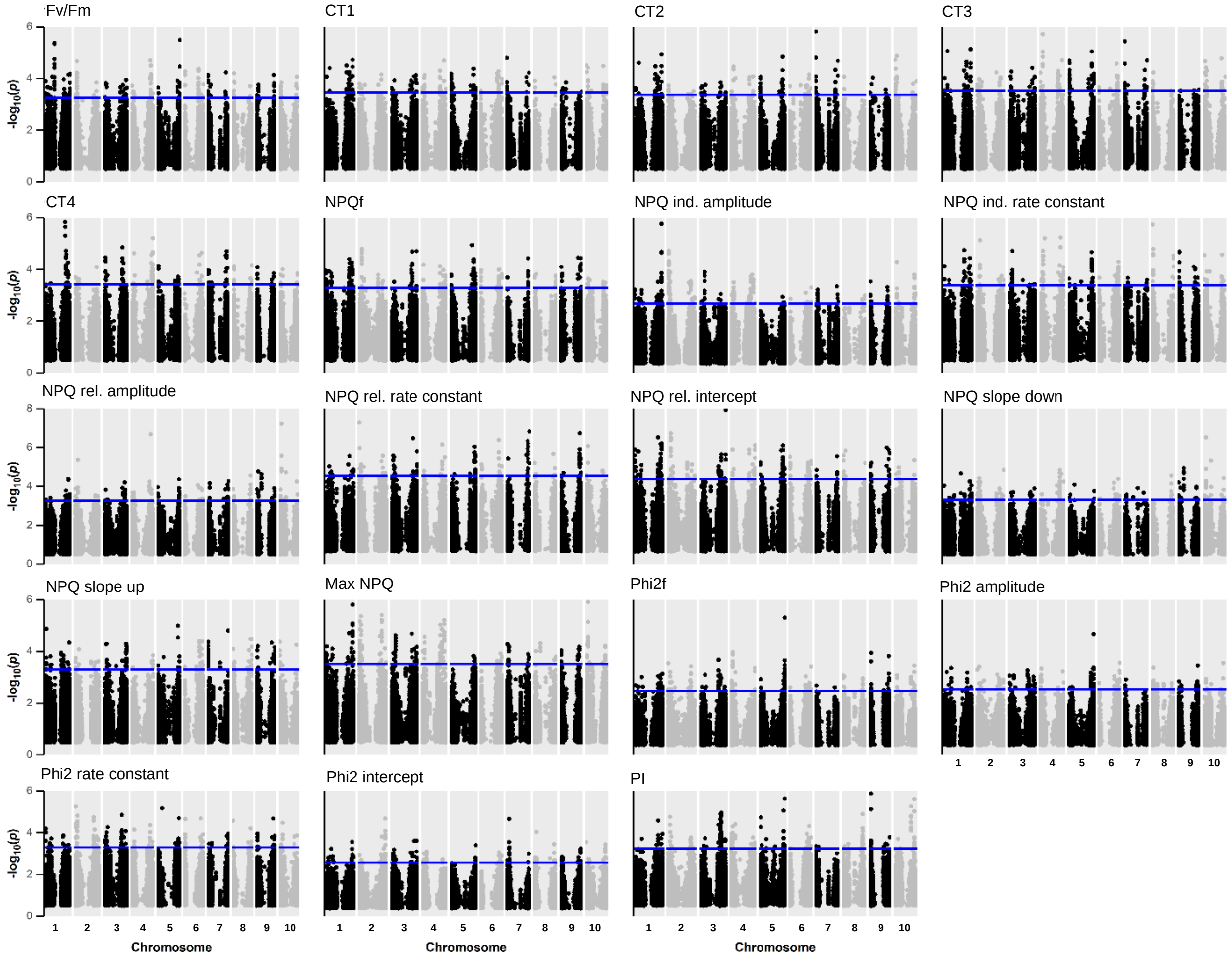


**Figure S13: Chromosome mapping (physical location) for genes associated with 2019 third-leaf tissue Fisher's combined test (FCT) non-photochemical quenching traits. Gene positions plotted as midpoint of each gene. Blue line indicates threshold of genes in top 1% by -log$_{10}$ *p*-value. CT: Combined trait analysis. CT1: Max NPQ, NPQ induction amplitude, NPQ induction rate constant *k*, NPQ relaxation rate constant *k*. CT2: NPQ induction amplitude, NPQ induction *k*, NPQ relaxation rate constant *k*. CT3: Max NPQ, NPQ induction rate constant *k*, NPQ relaxation rate constant *k*. CT4: $PI$, $\Phi PSII$ recovery amplitude, $\Phi PSII$ recovery rate constant *k*. NPQf: Final dark time point NPQ value. Phi2: Photosystem II quantum yield. Phi2f: Final dark time point Phi2. PI: Photoprotection index.**

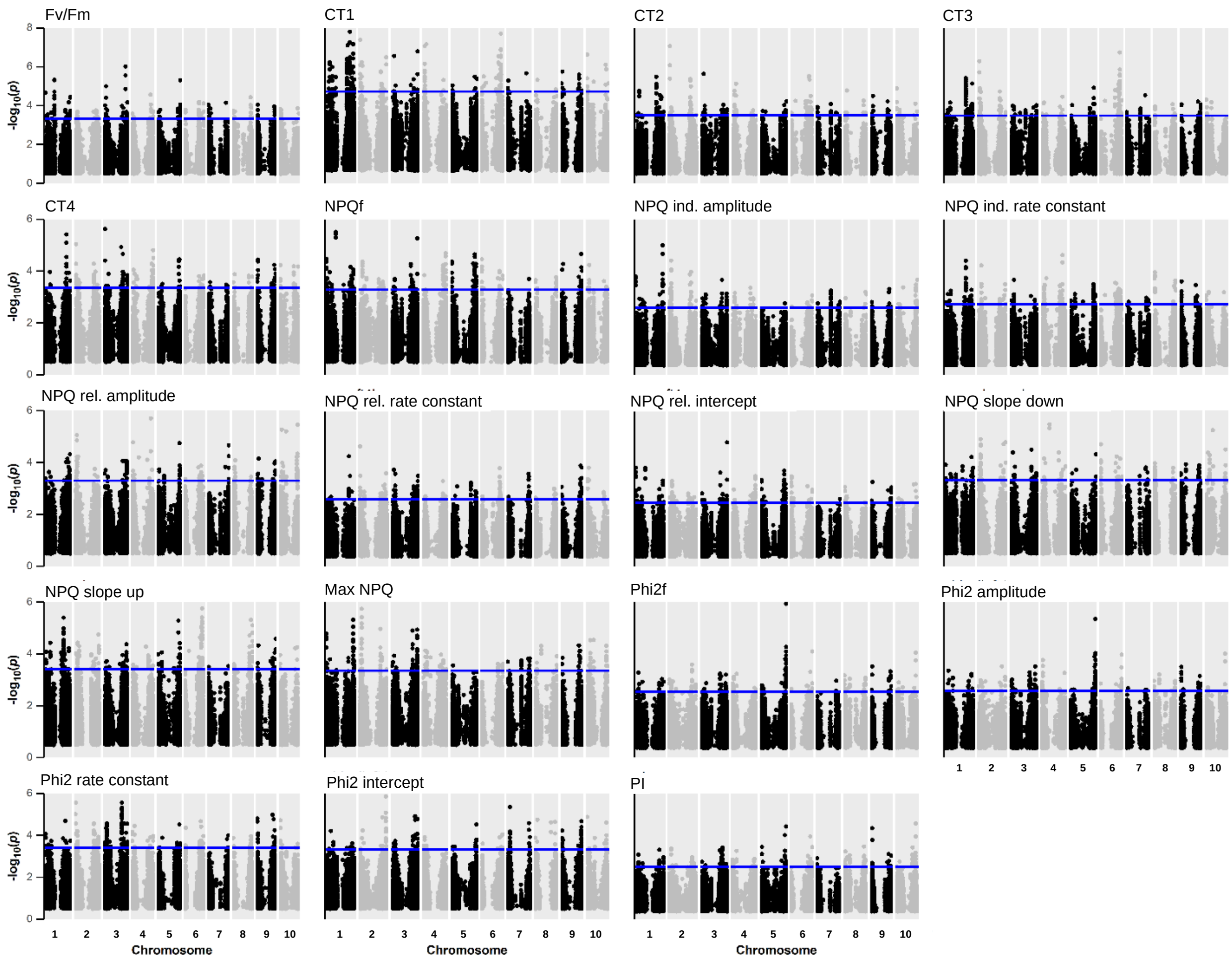


**Figure S14: Chromosome mapping (physical location) for genes associated with joint analysis third-leaf tissue Fisher's combined test (FCT) non-photochemical quenching traits. Gene positions plotted as midpoint of each gene. Blue line indicates threshold of genes in top 1% by -$\log_{10}$ *p*-value. CT: Combined trait analysis. CT1: Max NPQ, NPQ induction amplitude, NPQ induction rate constant *k*, NPQ relaxation rate constant *k*. CT2: NPQ induction amplitude, NPQ induction *k*, NPQ relaxation rate constant *k*. CT3: Max NPQ, NPQ induction rate constant *k*, NPQ relaxation rate constant *k*. CT4: $PI$, $\Phi PSII$ recovery amplitude, $\Phi PSII$ recovery rate constant *k*. NPQf: Final dark time point NPQ value. Phi2: Photosystem II quantum yield. Phi2f: Final dark time point Phi2. PI: Photoprotection index.**

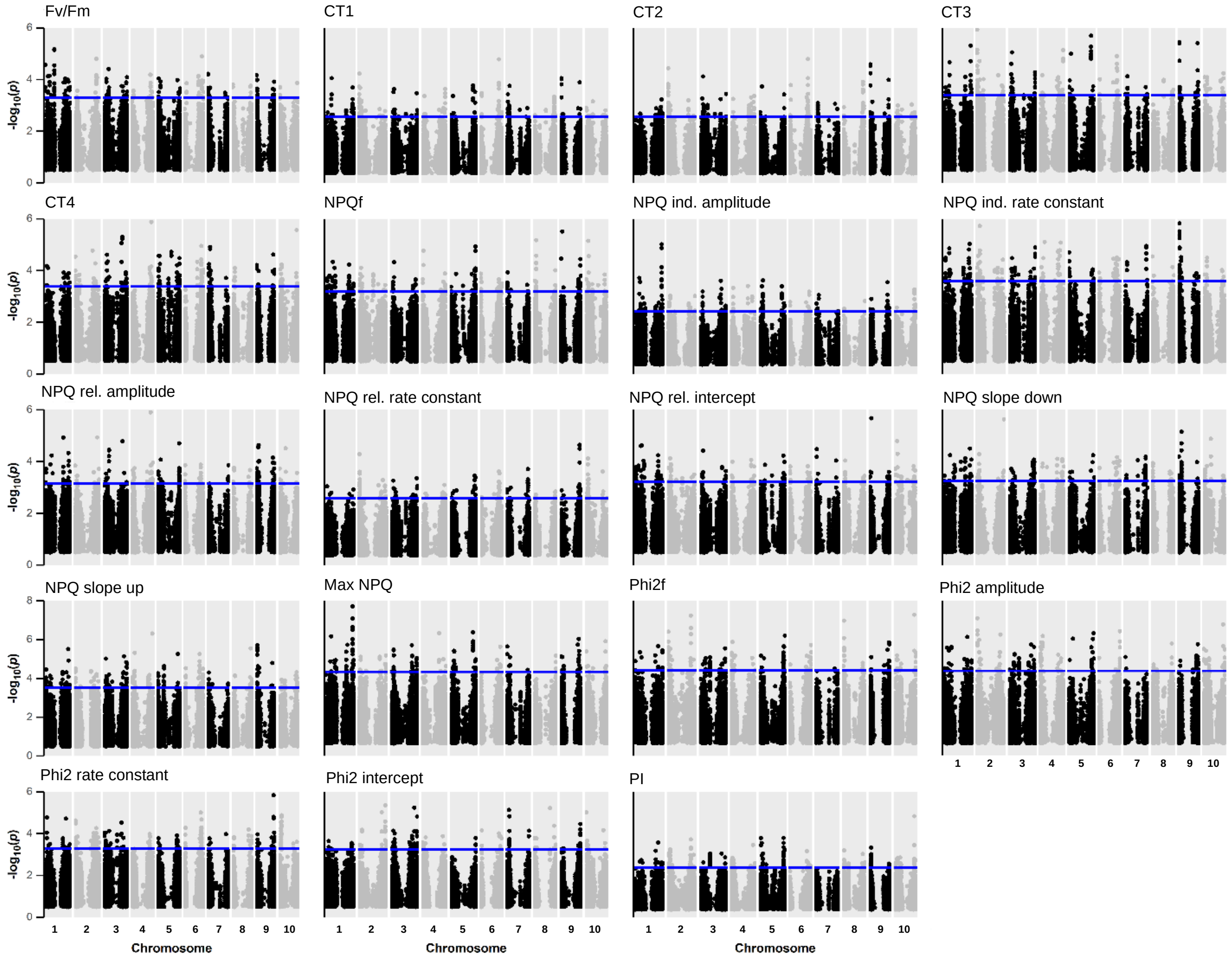


**Figure S15: Chromosome mapping (physical location) for genes associated with 2017 growing point tissue Fisher's combined test (FCT) non-photochemical quenching traits. Gene positions plotted as midpoint of each gene. Blue line indicates threshold of genes in top 1% by -log$_{10}$ *p*-value. CT: Combined trait analysis. CT1: Max NPQ, NPQ induction amplitude, NPQ induction rate constant *k*, NPQ relaxation rate constant *k*. CT2: NPQ induction amplitude, NPQ induction *k*, NPQ relaxation rate constant *k*. CT3: Max NPQ, NPQ induction rate constant *k*, NPQ relaxation rate constant *k*. CT4: $PI$, $\Phi PSII$ recovery amplitude, $\Phi PSII$ recovery rate constant *k*. NPQf: Final dark time point NPQ value. Phi2: Photosystem II quantum yield. Phi2f: Final dark time point Phi2. PI: Photoprotection index.**

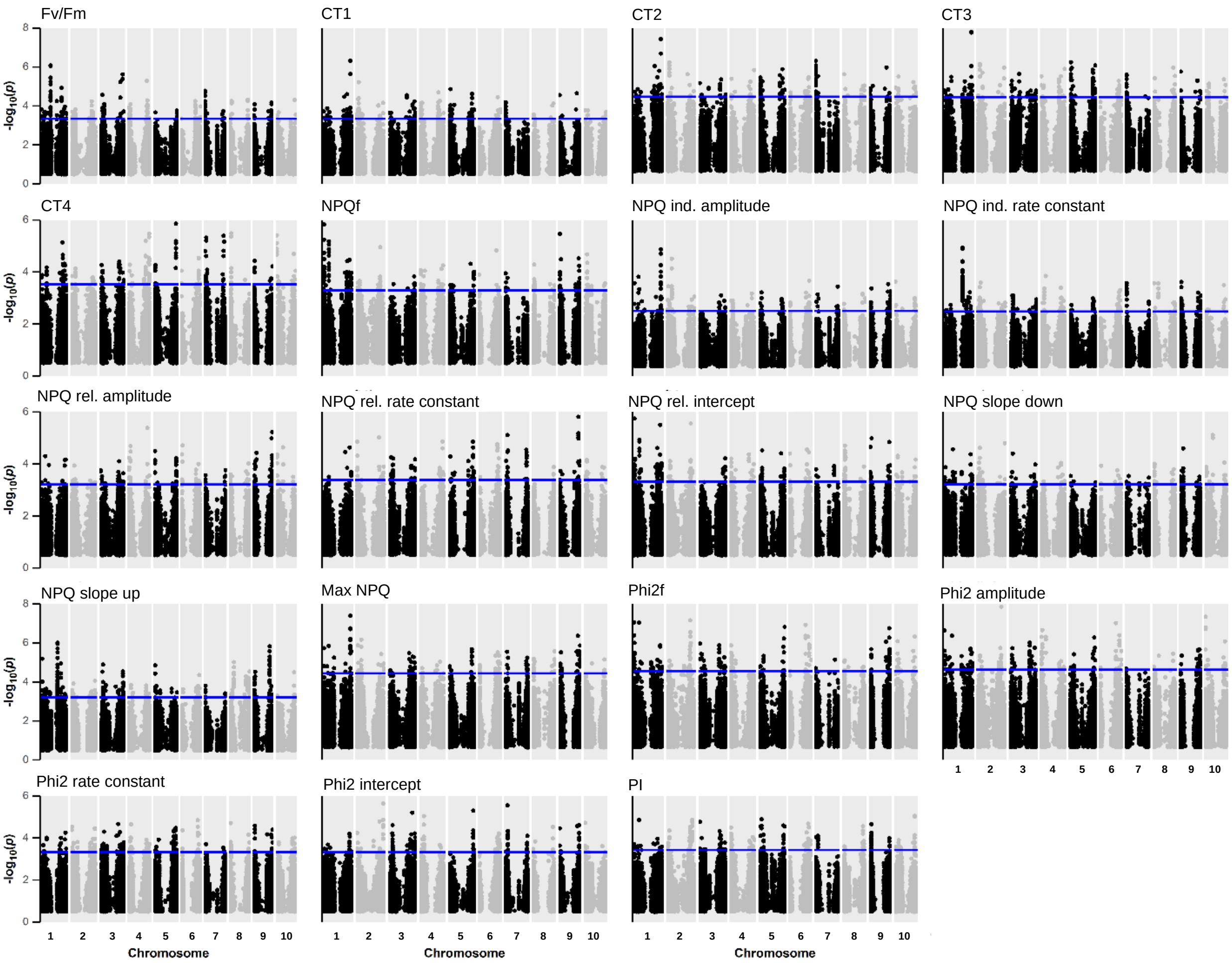


**Figure S16: Chromosome mapping (physical location) for genes associated with 2019 growing point tissue Fisher's combined test (FCT) non-photochemical quenching traits. Gene positions plotted as midpoint of each gene. Blue line indicates threshold of genes in top 1% by -$\log_{10}$ *p*-value. CT: Combined trait analysis. CT1: Max NPQ, NPQ induction amplitude, NPQ induction rate constant *k*, NPQ relaxation rate constant *k*. CT2: NPQ induction amplitude, NPQ induction *k*, NPQ relaxation rate constant *k*. CT3: Max NPQ, NPQ induction rate constant *k*, NPQ relaxation rate constant *k*. CT4: $PI$, $\Phi PSII$ recovery amplitude, $\Phi PSII$ recovery rate constant *k*. NPQf: Final dark time point NPQ value. Phi2: Photosystem II quantum yield. Phi2f: Final dark time point Phi2. PI: Photoprotection index.**

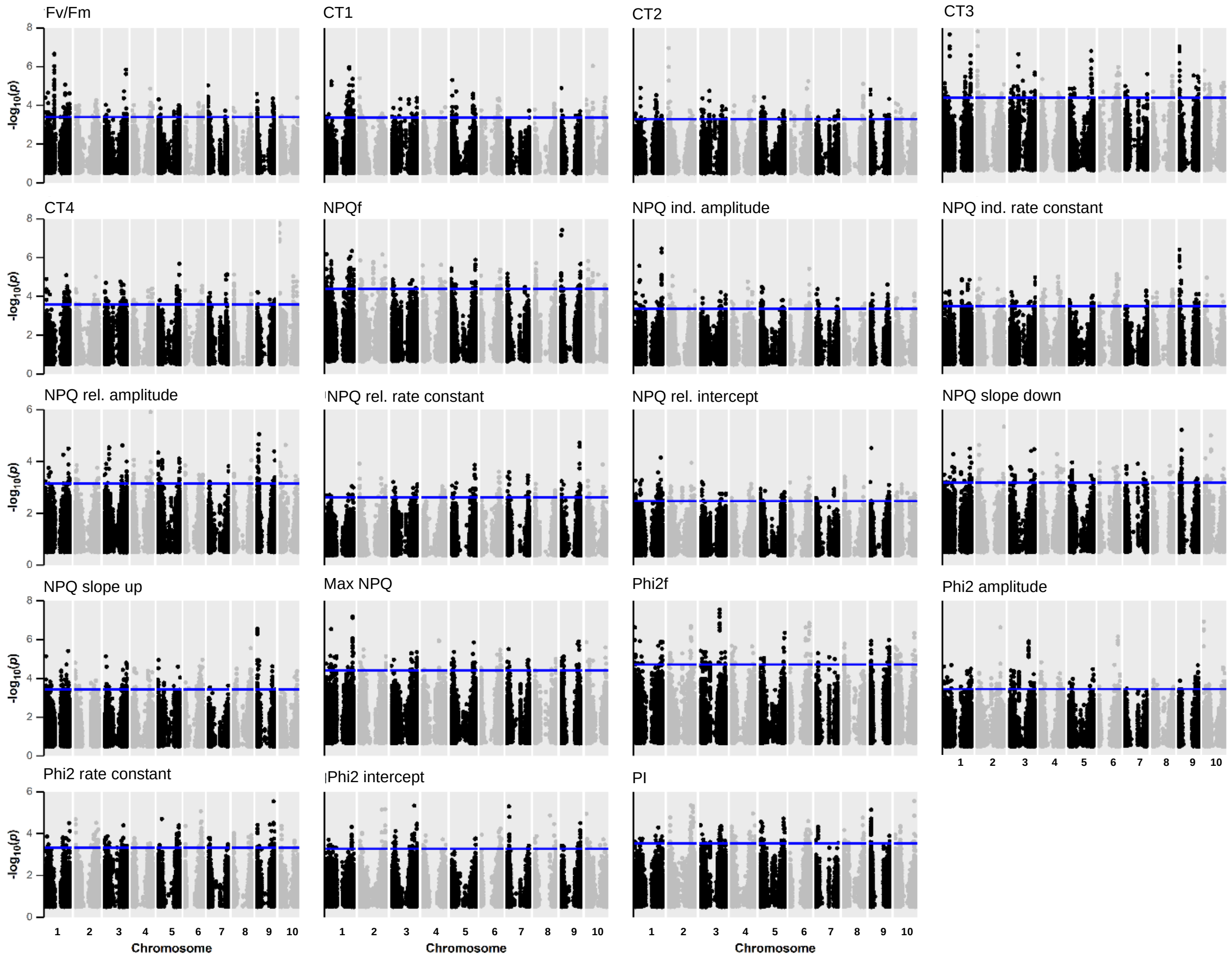


**Figure S17: Chromosome mapping (physical location) for genes associated with joint analysis growing point tissue Fisher's combined test (FCT) non-photochemical quenching traits. Gene positions plotted as midpoint of each gene. Blue line indicates threshold of genes in top 1% by $-\log_{10}$ *p*-value. CT: Combined trait analysis. CT1: Max NPQ, NPQ induction amplitude, NPQ induction rate constant *k*, NPQ relaxation rate constant *k*. CT2: NPQ induction amplitude, NPQ induction *k*, NPQ relaxation rate constant *k*. CT3: Max NPQ, NPQ induction rate constant *k*, NPQ relaxation rate constant *k*. CT4: *PI*, $\Phi PSII$ recovery amplitude, $\Phi PSII$ recovery rate constant *k*. NPQf: Final dark time point NPQ value. Phi2: Photosystem II quantum yield. Phi2f: Final dark time point Phi2. PI: Photoprotection index.**
